# Restoration of tumor suppressor protein with enhanced activity using super-tRNAs to induce tumor regression

**DOI:** 10.1101/2025.03.18.643282

**Authors:** Bangyan Kong, Zhetao Zheng, Zeyuan Mi, Zhiyang Dou, Yuelin Yang, Yuxuan Shen, Liangzhen Dong, Jingru Lv, Ying Li, Qing Liang, Songlin Tian, Jiayu Wang, Hua Qiao, Qing Xia

## Abstract

Cancer develops through a gradual accumulation of mutations including nonsense mutations in tumor suppressor genes (TSGs), which drive tumor proliferation and progression due to the impairment or complete loss of tumor suppressor protein function. The ability to restore functional tumor suppressor protein from mutant driver TSGs is a longstanding aspiration in cancer therapy, however, current strategies face formidable challenges in recovering proteins harboring heterogeneous mutations and circumventing dominant-negative effects exerted by endogenous mutant proteins. Therefore, the development of therapeutic interventions that are effective and generalizable across diverse nonsense mutations in cancer, to rescue endogenous mutant tumor suppressor proteins remains an urgent yet unmet need. Here we find that engineered suppressor transfer RNA (sup-tRNA) could overcome nonsense mutations in key driver tumor suppressor genes, such as *TP53*, thereby avoiding the dominant-negative effects through the restoration of endogenous mutant p53 with its original functions. Optimized sup-tRNA that we named super tRNA, enabling high-fidelity incorporation of desired amino acids, can further enhanced the transcriptional activity of p53 to suppress tumor cell proliferation. Moreover, we also establish a computational model to guide super tRNAs design, and demonstrate that super tRNA could effectively inhibit tumor growth by targeted readthrough of nonsense mutations in different tumor suppressor genes. In two xenograft models harboring nonsense mutant *TP53*, we observed up to a 70% reduction in tumor volume after lipid nanoparticle (LNP)-based sup-tRNA treatment, along with robust recovery of p53 expression and function, and no hematological or histological abnormalities detected. Our findings demonstrate the potential of sup-tRNA as a feasible and effective strategy to rescue and enhance tumor suppressor protein function, thereby preventing cancer development. As a novel therapeutic RNA, sup-tRNA provides a flexible strategy for targeting driver mutations in cancer, thereby expanding the potential of tRNA-based therapy in precision oncology.

## INTRODUCTION

Cancer is a disease of uncontrolled proliferation by transformed cells harboring mutations in driver genes including tumor suppressor genes and oncogenes^1–3^. Notably, over half of the high frequency mutations occur in tumor suppressor genes^4,5^, like *TP53*, mutations of which are found in approximately 42% of patients worldwide and their pivotal role in cancer development has been well established^6,7^. According to global cancer statistics reported by WHO, approximately 1 million new cancer cases annually harbor *TP53* nonsense mutations, leading to truncated inactive p53 proteins or complete loss of expression due to nonsense-mediated mRNA decay^8,9^ (NMD). However, rescuing these mutations remains a significant challenge in current therapies^10^. Unlike targeting mutations in oncogenes by inhibiting or degrading aberrant oncoproteins directly, rescuing tumor suppressor gene mutations requires the restoration of functional tumor suppressor proteins, which poses a formidable challenge to bind and recover mutant proteins of heterogeneous inactivating mechanisms with a single agent^11–13^. Large-scale studies have identified potential drug molecules, such as Rezatapopt and arsenic trioxide (ATO), which could stabilize p53 mutants and restore their functional conformation, through inserting into the Y220C crevice^14^ or filling the arsenic-binding pocket in p53 proteins, respectively^15–17^. However, their efficacy is clearly limited to p53 mutants with these specific binding sites and is not applicable for rescuing other *TP53* mutations, particularly nonsense mutations^11,18^.

As nonsense mutations result in premature termination of p53 translation, a facile and promising strategy is to restore endogenous full-length p53 by translational readthrough of nonsense codons during protein synthesis, avoiding undesirable adverse effects associated with exogenous p53 overexpression and the dominant-negative effects of truncated mutant proteins often observed in conventional gene therapy. In this context, developing therapeutic agents capable of recognizing and overriding only three nonsense codons (UAG, UAA, UGA) could address diverse *TP53* nonsense mutants, sparking renewed enthusiasm in recent years. To date, a few lead compounds, including aminoglycoside antibiotics and nucleoside analogs, have shown potential in overcoming cancer-derived *TP53* nonsense mutations and inhibiting tumor growth^19–22^. However, the reported readthrough approaches have yet to meet clinical needs, primarily due to their low efficiency (<10%) in most cases and the non-specific insertion of random amino acids in response to nonsense mutations, which may lead to the restoration of deleterious missense p53 mutants^23^.

Suppressor tRNA (sup-tRNA), derived from natural transfer RNAs, could recognize and pair with the nonsense codon during translation elongation, thus restoring the production and function of the full-length protein^24–26^. Given the high efficiency, low toxicity, and minimal safety concerns associated with sup-tRNA in treating inherited diseases caused by nonsense mutations, we anticipate that it also holds great promise for addressing nonsense mutations in human cancer. With this inspiration, we hypothesized that sup-tRNA can be used to restore p53 from nonsense mutations and induce tumor regression. Drawing from prior knowledge of p53 mutations, we further set out to established the framework to optimize sup-tRNA for rescuing diverse nonsense mutant p53 variants.

## RESULTS

### Engineering sup-tRNAs to target nonsense mutations in *TP53*

Tumor suppressor genes often harbor nonsense mutations, which cause abnormal termination of the translation of tumor suppressor proteins, thereby driving the development of tumors. For different tumor suppressor genes, the proportion of nonsense mutations in the total mutations varies greatly. We found that certain tumor suppressor genes (such as *TP53*, *RB1*, *PTEN*, and *APC*) often have a relatively high proportion of nonsense mutations. Furthermore, we quantify the “driver ability” of tumor suppressor genes according to the number of cancer types/cohorts in which this gene functions as a cancer driver gene. *TP53* is regarded as the gene with the strongest driver ability, as it exists as a mutational cancer driver gene in more than 50 types of cancer. Interestingly, our analysis of 315 driver tumor suppressor genes^2^ revealed a striking correlation: genes with higher driver ability exhibit a significantly higher proportion of nonsense/truncating mutations (*p* < 0.001), whereas synonymous mutations show an inverse trend (Figures 1A and S1A). This suggests that nonsense mutations, causing disastrous protein function loss, are evolutionarily selected in key tumor suppressors to promote tumor initiation (Figures 1B and S1-S2).

**Figure 1.**
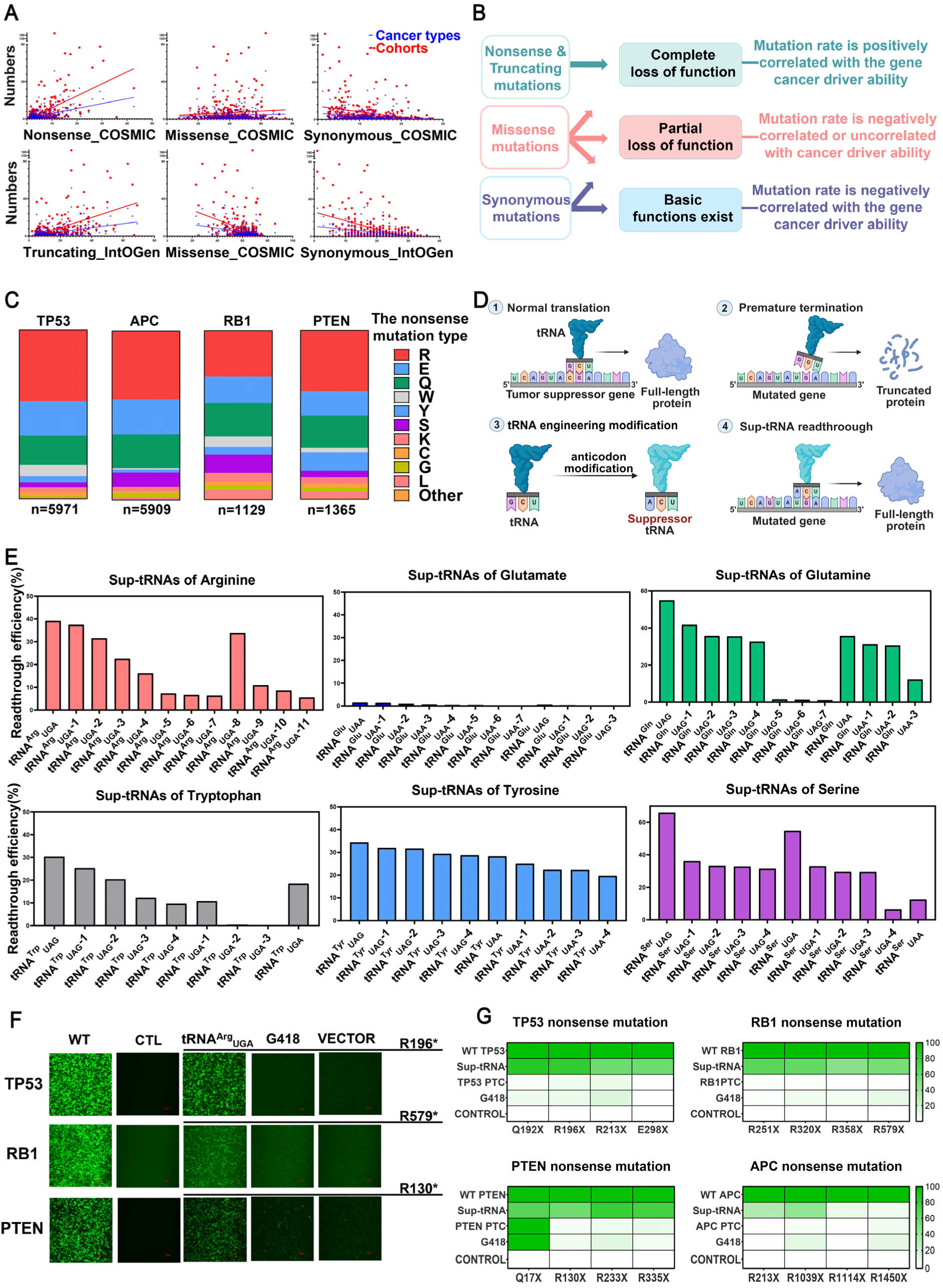
The efficacy of sup-tRNA in overcoming nonsense mutations within tumor suppressor genes. (A) Scatter plots of the proportion of mutation types (nonsense/ truncating mutation, missense mutation, synonymous mutation) and the number of cancer types and cohorts which specific tumor suppressor genes act as driver genes in the Catalogue of Somatic Mutations in Cancer (COSMIC) and Integrative OncoGenomics (IntOGen) for mutational cancer driver tumor suppressor genes. (B) The hypothesis explaining the correlation between the proportion of driver tumor suppressor genes mutation types and the cancer driver ability (In how many types of cancer/cohorts it serves as a driver gene). Based on the existing theories, nonsense mutation most leads to the complete loss of function thus missense mutation may cause complete loss of function or partial loss of function. The synonymous mutation leads to a partial loss of function or basic function. The variability in different mutation fates can rationally explain the correlation described above. (C) Classification of nonsense mutations according to their original codons and corresponding amino acids, and statistics of the mutation rates in tumor suppressor genes including *TP53*, *APC*, *RB1*, and *PTEN*, based on data adapted from COSMIC. (D) Schematics of engineering natural transfer RNA (tRNA) to suppressor tRNA (sup-tRNA) for nonsense mutations readthrough in tumor suppressor transcripts. (E) Quantification of readthrough efficiency of sup-tRNA encoding highly mutated amino acid defined in (C) using the dual-luciferase reporter assay. (F and G) Microscopy images (F) and quantification (G) of GFP positive cells showing differential readthrough efficiencies of above three approaches. Consistent results were obtained independently for at least three biological replicates. In (F and G), all data shown from HEK 293T cell experiments are mean of three biological replicates. Scale bar=100 µm.

We then analyzed the types of amino acids affected by nonsense mutations in the four vital tumor suppressor genes *TP53*, *APC*, *RB1*, and *PTEN*. The amino acids involved in these mutations were similar across these genes, primarily consisting of arginine (Arg), glutamic acid (Glu), and glutamine (Gln) (Figure 1C). With the exception of glutamic acid, all frequently mutated amino acids had corresponding suppressor tRNAs capable of inducing nonsense readthrough by up to 55%, offering potential for the restoration of tumor suppressor proteins (Figures 1D-1E). Next, we constructed a series of surrogate reporters harboring nonsense mutant tumor suppressor genes and EGFP genes linked by a 2A peptide sequence, and employed sup-tRNAs to induce targeted readthrough of nonsense codons (UAG, UGA) to permit the downstream enhanced green fluorescent protein (EGFP) expression, allowing us to measure the readthrough efficiency through monitoring EGFP level. Though readthrough efficiency varies among different genes, we found that sup-tRNA consistently outperforms the documented readthrough-promoting agent G418 (geneticin), achieving up to a 276-folds increase in efficiency (Figures 1F-1G). These results indicate that sup-tRNA can efficiently overcome nonsense mutations in tumor suppressor genes, inducing nearly 70% readthrough in nonsense mutant *TP53*. The actual amino acids incorporated by these sup-tRNAs have been verified by mass spectrometry. These results suggested that sup-tRNA enabled the efficient and precise incorporation of specific amino acids at nonsense mutations.

To further investigate the capability of sup-tRNAs for the functional restoration of p53, we co-transfected NCI-H1299 cells (*TP53*-null) with sup-tRNAs, the p53 reporter, and a genetically encoded sensor, which allowed us to characterize full-length p53 protein expression via green fluorescence and assess its transcriptional activity restoration through red fluorescence (Figure 2A). We found that sup-tRNAs carrying glutamine or arginine were able to rescue the expression and function of the frequent p53 mutants, albeit with varying efficiencies (Figures 2B-2C and S3-S5). Furthermore, functional restoration exhibited a significant correlation with readthrough efficiency. However, when applying the high-efficiency sup-tRNAs to address other COSMIC-reported p53 mutants harboring UAG or UGA nonsense mutations, we observed that the function of two mutants (G266* and Y234*) was not restored, despite the recovery of full-length protein expression (Figures 2B and S4). By analyzing Y234* and G266* variants based on p53 crystal structure with molecular dynamics simulation, we speculated that mischarging of Y234Q and G266R in p53, induced by sup-tRNA^Gln^_UAG_ and sup-tRNA^Arg^_UGA_, may be deleterious (Figure 2E). This is primarily attributed to the disruption of a hydrophobic core around Y234 and the formation of new hydrogen bonds between G266 and adjacent residues, potentially compromising the p53-DNA binding interface (Figure 2E). Therefore, a reliable approach for restoring p53 activity necessitates the simultaneous consideration of readthrough efficiency and the functional impact of the incorporated amino acid.

**Figure 2.**
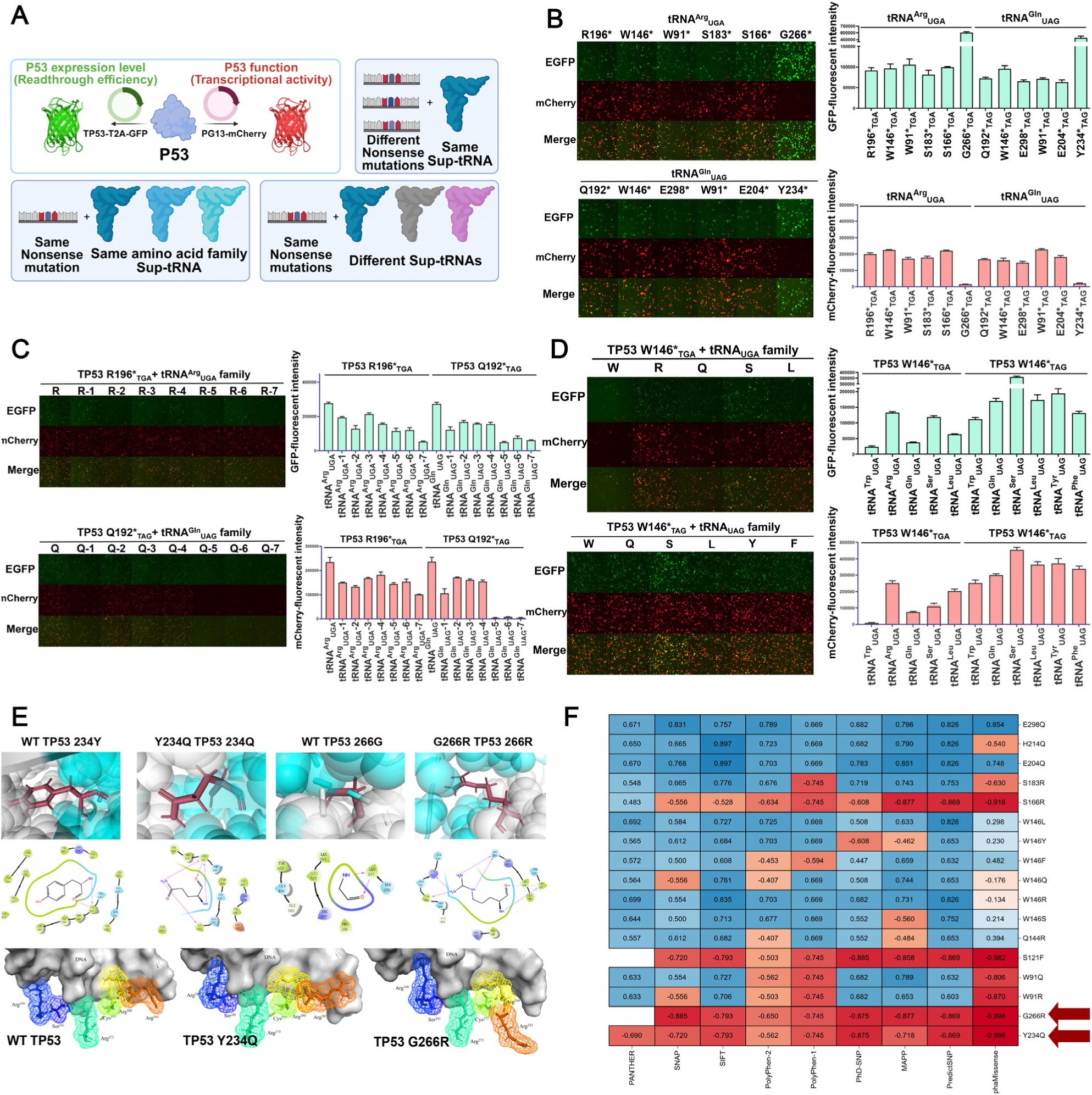
Characterization of sup-tRNA in recovering p53 expression and function. (A) Schematic of a genetically encoded reporter system to measure the expression and function of p53 via dual-fluorescence detection, which is designed for sup-tRNA efficiency characterization. (B) Representative microscopy images (left) and quantification of GFP and mCherry positive cells (right) show p53 expression and function in NCI-H1299 cells with 12 kinds of nonsense mutations after tRNA^Arg^_UGA_ and tRNA^Gln^_UAG_ readthrough. Data are mean ± s.d. of three biological replicates. Scale bars = 100 µm. (C) Representative microscopy images (left) and quantification of GFP and mCherry positive cells (right) show p53 expression and function in NCI-H1299 cells with *TP53* R196*& *TP53* Q192* nonsense mutations after 8 kinds of arginine (R) and 8 kinds of glutamine (Q) sup-tRNAs readthrough. Data are mean ± s.d. of three biological replicates. Scale bars = 100 µm. (D) Representative microscopy images (left) and quantification of GFP and mCherry positive cells (right) show p53 expression and function in NCI-H1299 cells with *TP53* W146* nonsense mutations after 7 amino acids while 12 kinds of high-efficiency sup-tRNAs readthrough. Data are mean of three biological replicates. Scale bars = 100 µm. (E) Verified deleterious p53 variants mutant site amino acid residues and intermolecular forces. The first and third panels show the 234Y, and 266G structures of wild-type (WT) p53 DBD (2AC0), respectively. The other panels are drawn by mutating 234Y, 266G residue in wild-type DBD (2AC0) into the indicated residues in PyMOL and Schröder. The panels below show the close-up view of the wild-type and verified deleterious p53 DNA binding peptides and DNA. The side chains of the six classic DNA binding residues are shown as sticks. (F) Predictive and functional profile of p53 previous variants. The heat map corresponds to the predictive impact of each p53 variant ranging from -1 (red) to 1 (blue), with the lowest score being the most deleterious. Scores from the 9 algorithms in the panels including AlphaMissense, MAPP, PANTHER, PhD-SNP, PolyPhen-1, PolyPhen-2, SIFT, and SNAP. Red arrows indicate previous verified deleterious p53 variants.

### Computationally optimized screening of sup-tRNAs for enhanced p53 activity restoration

To this end, we developed a computational tool^27,28^ to predict the functional impact of incorporated amino acids, facilitating the selection of optimal sup-tRNAs (Figure 2F). Using the p53-E204* mutant cancer cell line containing a TAG codon, we assessed its expression and functional restoration following readthrough induced by sup-tRNAs carrying different amino acids (Figure S7A). Consistent with the prediction, we found that tRNA^Tyr^_UGA_, tRNA^Phe^_UGA_, and tRNA^Trp^_UGA_ failed to restore p53 transactivation function due to the insertion of undesired amino acids, despite its high readthrough efficiencies (Figure S7B). Moreover, even when the original amino acid could not be restored due to the low readthrough efficiency of the corresponding tRNA^Glu^_UGA_, high-efficiency sup-tRNAs, such as tRNA^Gln^_UGA_ and tRNA^Ser^_UGA_, served as effective alternatives to rescue p53 function, provided that the incorporated amino acid, as predicted, does not disrupt the protein structure (Figure S8-S9). Further investigation also confirmed the robustness of this alternative strategy in restoring other functional p53 mutants, however, we noticed that some recovered p53 variants showed superior transcriptional activity compared to wild-type p53 proteins, for instance, tRNA^Arg^_UGA_ outperformed tRNA^Trp^_UGA_ in functional restoration of p53-W146*_TGA_ mutant (Figure 2D), even though the latter enabled recovery of wild-type p53 proteins.

These observations can be mechanistically explained by two critical determinants governing p53 functional restoration. First, when the incorporated amino acid induces minimal structural perturbation to native conformation of p53, sup-tRNAs with higher readthrough efficiency, such as tRNA^Gln^_UAG_, sup-tRNA^Arg^_UGA_, can achieve superior functional restoration (Figures 2B-2D and S4-S6). Second, in cases where readthrough efficiencies are comparable across different sup-tRNAs, the identity of the restored amino acid becomes particularly crucial.

### Harnessing sup-tRNAs to overcome endogenous *TP53* nonsense mutations

Genomic analyses from COSMIC and intOGen databases have consistently identified *TP53* nonsense mutations as one of the most prevalent genetic alterations in non-small cell lung carcinoma (NSCLC) and colorectal cancer (CRC) (Figures S10A-S10B). TCGA data further highlight *TP53* as the most critical driver gene in both NSCLC and CRC (Figures S10C-S10D). Therefore, we further investigated the efficacy of sup-tRNAs using NSCLC cell line Calu-6 (*TP53* R196*_TGA_) and CRC cell line Caco-2 (*TP53* E204*_TAG_), whose nonsense mutant *TP53* genes have been verified by whole-genome sequencing (Figures S10E-S10F). In Calu-6 cells, tRNA^Arg^_UGA_, engineered from an arginine-carrying tRNA with a UCU anticodon, effectively restored high levels of full-length p53 expression (Figure 3A). In contrast, G418 treatment resulted in only minimal p53 recovery, comparable to the tRNA^Arg^_UGA_ control and even significantly lower than that achieved with the low-efficiency (∼6%) tRNA^Arg^_UGA_-7 (Figure 3A). In Caco-2 cells, tRNA^Gln^_UAG_ also yielded a robust restoration of full-length p53 protein expression compared to other groups, as determined by western blot analysis (Figure 3A). The recovery of p53 transactivation function was confirmed by red fluorescent encoded by the sensor (Figures 3C and S11) and increased expression of cyclin-dependent kinase inhibitor 1A (CDKN1A), a major downstream target of p53 activity, and the recovery of p53 expression further validated through immunostaining (Figures 3A-3B).

**Figure 3.**
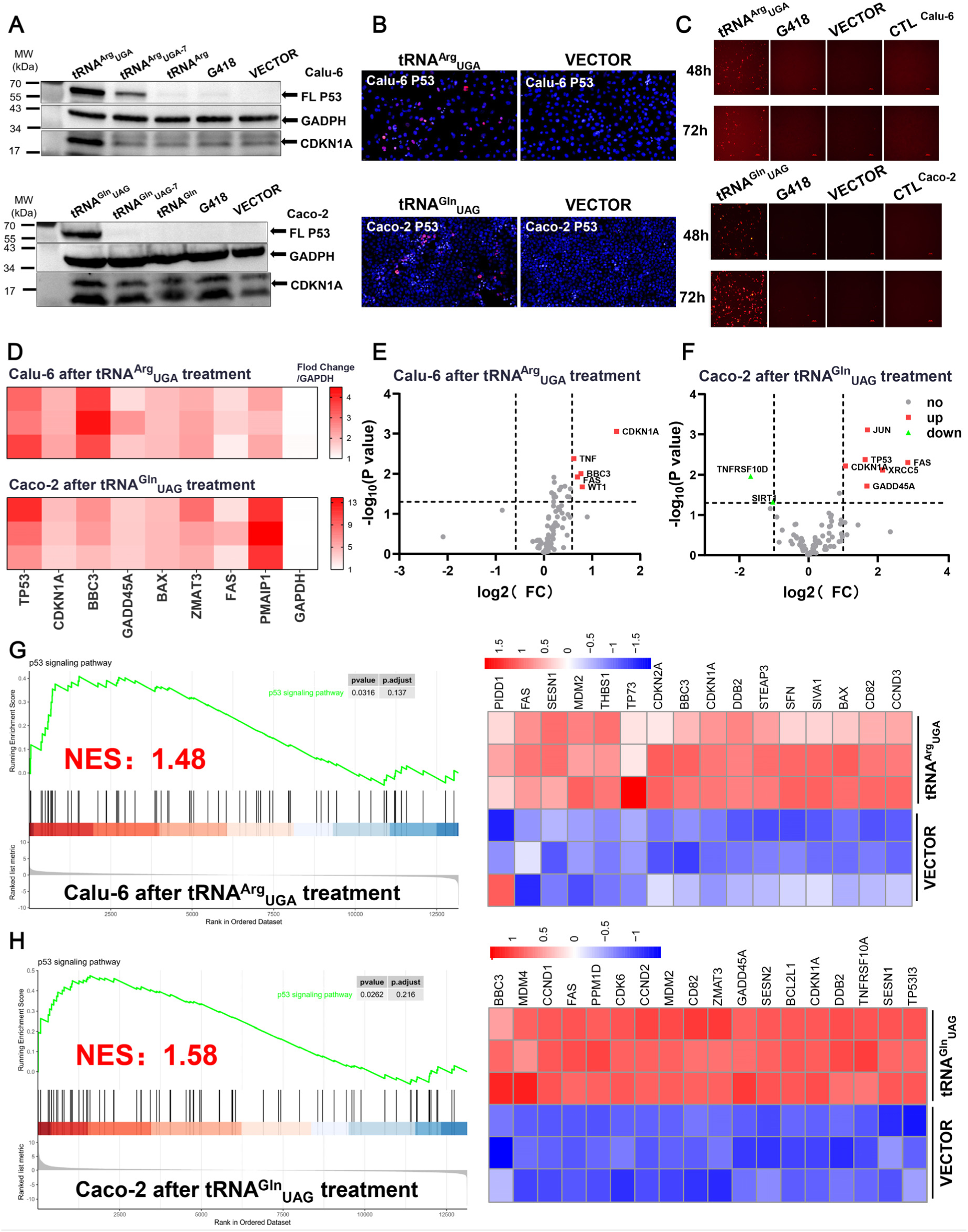
Sup-tRNA rescues endogenous full-length p53 expression and restores its transcriptional activity. (A) Western blot analysis of full-length p53 recovery and protein expression from its target gene *CDKN1A* in Calu-6 (above) & Caco-2 cells (below) treated with sup-tRNA, corresponding tRNA, vector, and G418 for 48 hours, respectively. (B) Representative microscopy images of immunofluorescence for full-length p53 and CDKN1A in Calu-6 (above) & Caco-2 cells (below) treated with sup-tRNA or vector for 48h. Scale bars= 20 µm. Data shown here were representative examples from n=3 independent biological replicates. (C) Representative microscopy images showing the transcriptional activity of endogenous p53 in Calu-6 (above) & Caco-2 cells (below). treated with sup-tRNA, corresponding tRNA, vector, and G418 for 48&72 hours Data shown here were representative examples from n=3 independent biological replicates. (D) Quantification of mRNA levels of *TP53* and its target genes *CDKN1A*, *BBC3*, *GADD45A*, *BAX*, *ZMAT3*, *FAS*, and *PMAIP1* after sup-tRNA treatment in Calu-6 (above) & Caco-2 cells (below) for 48h by qRT-PCR. All the mRNA expression levels are normalized to *GAPDH*. Data are represented as mean for n = 3 independent biological replicates. (E and F) Significant differential expressed of p53-relatived transcripts are determined by p53 PCR array on Calu-6 (E) & Caco-2 (F) treated with sup-tRNA, or vector control, n = 3. (G and H) Gene Set Enrichment Analysis (left) and gene expression analysis (right) of p53 signaling pathways on Calu-6 (G) & Caco-2 cells (H) treated with sup-tRNA, or vector control, n = 3.

Classically, p53 functions mainly as a sequence-specific transcription factor that can bind to p53 response element and activate the transcription of target genes. To evaluate the transcriptional activity of restored p53, we examined the expression levels of its target genes using RT-qPCR. Our results showed that the RNA expression levels of p53 and its downstream targets were upregulated to varying degrees in both Calu-6 and Caco-2 cell lines (Figure 3D). Notably, the extent of upregulation increased over time, suggesting that sup-tRNA may prevent *TP53* mRNA with nonsense mutations from degradation by nonsense-mediated mRNA decay^9^ (NMD), thereby promoting full-length p53 production for transactivation. To further confirm the efficacy of sup-tRNA in reactivating p53-related genes, we performed a PCR array analysis. In Calu-6 cells, RNA expression levels of p53 target genes, including *CDKN1A* and *BBC3*, were differentially upregulated following readthrough restoration of p53 (Figure 3E). Similarly, in Caco-2 cells, RNA expression levels of *TP53* and its downstream targets were differentially upregulated, which consistent with RT-qPCR results (Figure 3F).

Moreover, RNA-seq analysis enabled us to assess the global influence of sup-tRNA on the transcriptomes of Caco-2 and Calu-6 cells. Gene Set Enrichment Analysis (GSEA) revealed that, compared with the control group, sup-tRNA treatment significantly upregulated the p53 signaling pathway in tumor cells from both Calu-6 (NES = 1.48, *p* = 0.0316) and Caco-2 (NES = 1.58, *p* = 0.0262) (Figures 3G-3H). The mRNA expression of p53-target genes also showed a significant increase compared to the control group (Figures 3G-3H).

### Suppressing tumor cell proliferation by restoring p53 function

To verify that sup-tRNA could inhibit tumor cell growth by suppressing nonsense mutations in the *TP53* gene, we transfected *TP53*-null NCI-H1299 cells with both a nonsense mutant *TP53* and sup-tRNA, or with either component individually (Figure 4A). The colony formation assay demonstrated that sup-tRNA exerted tumor-suppressive effects only in the presence of *TP53* nonsense mutations readthrough, leading to approximately a 60% reduction in colony number, a 70% decrease in total colony area, and a 30% reduction in the average area of individual clones (Figure 4B). In contrast, sup-tRNA alone did not exhibit significant proliferation-inhibitory effects in tumor cell lines lacking nonsense mutant *TP53* (Figure 4B).

**Figure 4.**
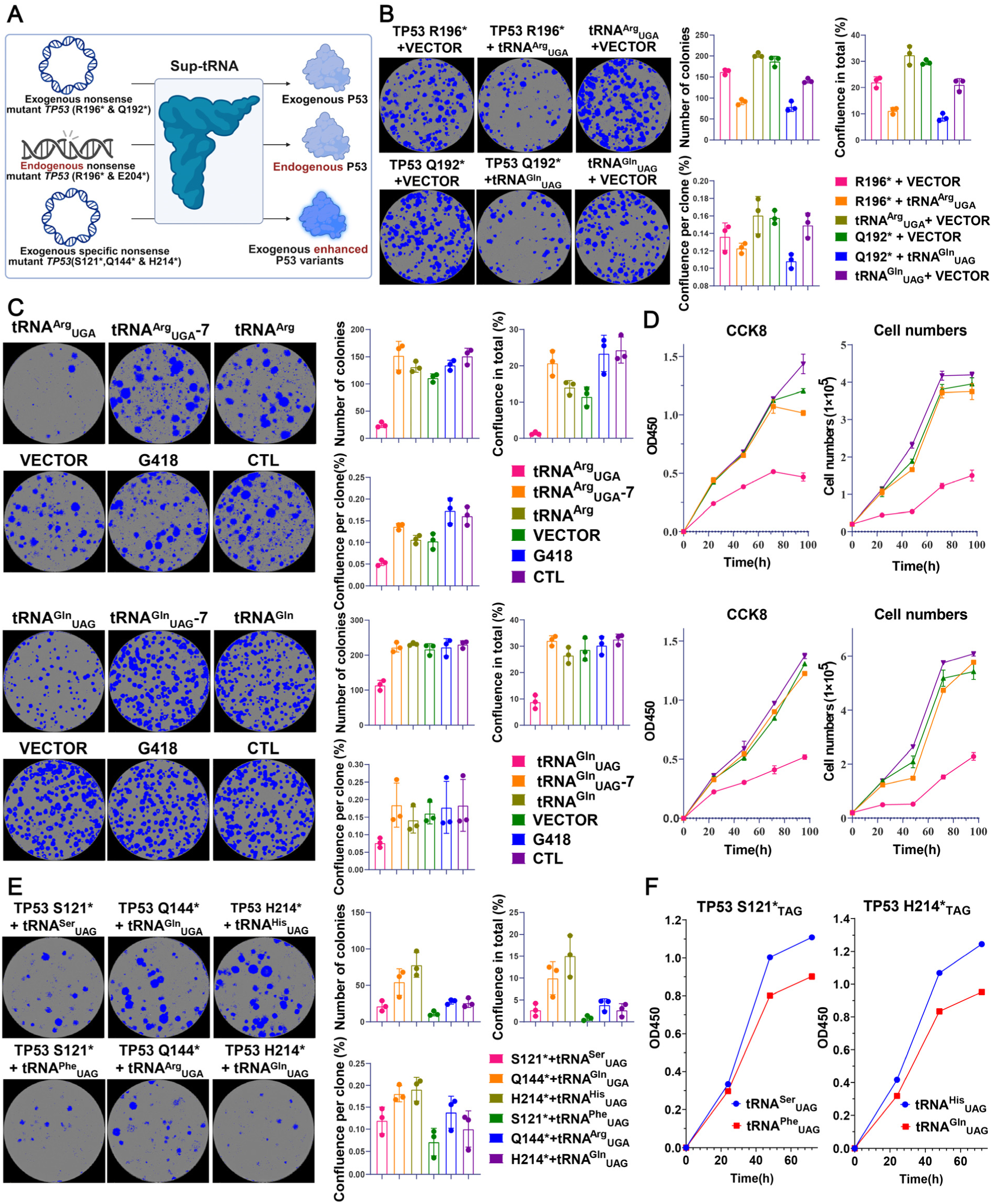
Sup-tRNA elicits antiproliferation activity through p53 restoration. (A) Schematic workflow for measure antiproliferative effects mediated by p53 restoration via sup-tRNA readthrough. (B) Sup-tRNAs in the presence of *TP53* exogenous nonsense mutation reduces the colony formation in human NCI-H1299 cells. Colonies are stained with crystal violet staining solution and counted after 10-12 days of culture. Colony images are acquired by Incucyte SX5 and processed and analyzed by Incucyte 2022B Rev2. Data are represented as mean ± SEM, n = 3. (C) Sup-tRNA treatment reduce the colony formation in human Calu-6 (above) & Caco-2 (below) cell which have genomic *TP53* nonsense mutation. Colonies are stained with crystal violet staining solution and counted after 10-12 days of culture. Colony images are acquired by Incucyte SX5 and processed and analyzed by Incucyte 2022B Rev2. Data are represented as mean ± SEM, n = 3. (D) Cell proliferation curve was measured by CCK-8 (left) and cell count assay (right). After transfecting human Calu-6 (above) & Caco-2 (below) cell for 8 hours, the cells are digested, spread in plates and further measured. Data are represented as mean ± SEM, n = 3. (E) The superb-tRNA strategy compared to the strategy restore as wild-type p53 reduces the colony formation in human NCI-H1299 cell which introduce exogenous *TP53* nonsense mutation. Colony images are acquired by Incucyte SX5 and processed and analyzed by Incucyte 2022B Rev2. Data are represented as mean ± SEM. N = 3. (F) CCK8 shows the superb-tRNA strategy compared to the strategy restore as wild-type p53 had a stronger growth inhibitory effect on the human NCI-H1299 cell which introduces exogenous *TP53* nonsense mutation.

Next, we sought to evaluate the efficacy of sup-tRNA in suppressing the proliferation of tumor cells harboring endogenous *TP53* nonsense mutations. Treatment with high-efficiency sup-tRNAs resulted in significant inhibition of cell growth and reduced cell viability. In Calu-6 cells, restoration of p53 using tRNA^Arg^_UGA_ resulted in a notable reduction in colony number, total colony area, and individual clone size, outperforming both G418 treatment and the low-efficiency control, consistent with previous findings (Figures 4C-4D). Similar results were also observed in Caco-2 cells, further highlighting the robustness of sup-tRNA across different cell lines (Figures 4C-4D). Given that nonsense mutant *TP53* could produce missense p53 variants with enhanced tumor suppressor function^29^ via sup-tRNA-mediated readthrough (Figure 4A), we further demonstrated the augmented efficacy of restored p53 variants S121F, Q144R, H214Q in inhibiting tumor cell growth (Figures 4E-F and S12-S13). As conventional gene therapy could only introduce wild-type p53 that may be neutralized by endogenous truncated p53 in a dominant-negative effect^30^, high-efficiency sup-tRNAs could directly convert truncated p53 into functional p53 variants^29^(Figure S14) in suppressing tumor cell proliferation, providing a plausible strategy for cancer therapy.

### Super- and superb-tRNA strategies for tRNA therapy

By applying first principles thinking, we have delineated tRNA therapy into two determinants: (1) Efficient restoration of full-length protein expression, which is governed by the readthrough efficiency of the specific sup-tRNA. (2) Functional recovery of restored protein, influenced by properties of the incorporated amino acid at nonsense mutations. Therefore, the most straightforward strategy for tRNA therapy is to restore original amino acids at nonsense mutations, which is particularly applicable to arginine and glutamine mutations, due to the high efficiency of tRNA^Arg^_UGA_ and tRNA^Gln^_UAG_ (Figure 1E). However, this strategy cannot facilitate the readthrough of approximately 43% nonsense mutations in *TP53*, since the low efficiency of corresponding sup-tRNAs. Thus, we reasoned that an alternative is using high efficiency sup-tRNA to convert the nonsense mutation to a missense but neutral mutation in proteins, based on computational analysis, which is named as super-tRNA therapy (Figure S15A). Meanwhile, we defined the strategy of converting nonsense mutations into wild-type proteins through high efficiency sup-tRNAs, which was previously used in treating inherited diseases, as primary-sup-tRNA therapy. Conversely, utilizing sup-tRNAs to restore deficient protein expression, or to cause missense and deleterious mutations, called lower-sup-tRNA therapy, ought to be avoided in future applications (Figure S15A).

Aforementioned results suggested the feasibility of super-tRNA in multiple aspects. For *TP53* W146*_TGA,_ although both tRNA^Trp^_UGA_ and tRNA^Arg^_UGA_ could suppress this nonsense mutation, tRNA^Arg^_UGA_, increased the restored protein expression level by 5.7 folds and the transcriptional function by 27 folds, compared with tRNA^Trp^_UGA_. Similarly, for *TP53* W146*_TAG_, tRNA^Ser^outperformed tRNA^Trp^_UAG_, in increasing the restored protein expression level by 3.2 folds and the transcriptional function by 1.8 folds (Figure 2D). It is worth noting that super-tRNA is engineered for restoring nonsense mutations to neutral mutations, but not for restoring them to deleterious missense mutations. For example, we attempted to restore *TP53* G266*_TGA_ and Y234*_TAG_ using tRNA^Arg^_UGA_ and tRNA^Gln^_UAG_ respectively. The efficient restoration of protein expression was achieved by both sup-tRNAs, but neither could rescue protein functions. We attribute this to the restoration of the *TP53* G266R and Y234Q mutations, which are considered deleterious mutations (Figures 2B and S4). Surprisingly, all these deleterious mutations can be accurately identified by the computational tools, which indicates that super-tRNA therapy can be safely applied with the assistance of the computational tools. These results suggest that a single super-tRNA could be employed to treat 10 different amino acid types of nonsense mutations, which would significantly reduce the cost in developing drug development (Figures 2F and S15B).

In addition to recovery of wild-type or neutral p53 variants, we anticipated that sup-tRNA could produce p53 with enhanced functions, via high-fidelity specific amino acid incorporation at nonsense mutations (Figure S15A). Some p53 variants, including S121F, H214Q, and Q144R, have been shown to exhibit enhanced apoptotic induction compared to wild-type p53, suggesting that enhancing p53 function would contribute to tumor inhibition ^29^. Here we termed this sup-tRNA based strategy as superb-tRNA strategy (Figure 15A) and attempted to restore *TP53* S121*, H214*, and Q144* to produce advantageous p53 variants with enhanced anti-cancer function (Figure 4A). The superb-tRNA strategy achieved more efficient tumor suppression effects at all the three aforementioned nonsense mutations. Compared with restoration of wild-type p53, all advantageous p53 variants achieved more pronounced tumor suppression, resulting in up to 80% reduction in colony number and 90% reduction in total colony area (Figures 4E-4F), which is consistent with previous reports^29^.

Therefore, by leveraging the capacity of tRNAs to precisely incorporate desired amino acids during protein translation, we could restore p53 variants with retained (super-tRNA therapy) or even enhanced functions (superb-tRNA strategy), supported by computational guided sup-tRNA selection (Figures 2F and S15). We also noted that a recent investigation employed CRISPR-mediated homology-directed repair to engineer 9,225 p53 variants in cancer cells, covering over 94% of cancer-associated missense mutations, and mapped their effects on p53 function^31^. These precious data are also helpful for choosing specific sup-tRNA for super-tRNA and superb-tRNA therapy .

### *In vivo* efficacy of sup-tRNA in *TP53* nonsense mutant xenograft

Lipid nanoparticle (LNP)-based RNA therapy has emerged as a pioneering platform in modern drug development. Its success in the COVID-19 pandemic has validated its potential for rapid, scalable, and adaptable medical solutions. As a novel therapeutic RNA, sup-tRNA can in theory be delivered to tumors via LNP. Here we utilized classical FDA-approved LNP formulation in which SM-102 serving as the ionizable lipid. All LNP-tRNAs (including tRNA^Arg^_UGA_ & tRNA^Gln^_UAG_) formulations were monodisperse spherical nanoparticles with a size of approximately 150 nm and possessed a slightly negative zeta potential (Figure S16A-S16B), indicating that LNP-tRNA have a generally uniform particle size distribution suitable for cellular uptake and endosomal escape, thus facilitating cancer treatment *in vivo*.

We next utilized cell-line-derived xenograft (CDX) mouse models to investigate whether sup-tRNA could restore p53 protein expression from *TP53* nonsense mutations *in vivo* and inhibit tumor growth. Six-week-old immunodeficient nude mice were implanted with either Calu-6 or Caco-2 cells to generate xenograft models harboring *TP53* nonsense mutations, and were treated with LNP-tRNA via intra-tumoral delivery once the tumor volume reached approximately 200 mm³ (Figure 5A). For the NSCLC Calu-6 xenograft model with p53-R196*, we treated the mice with LNP-tRNA^Arg^_UGA_ and G418 as a positive control. The tumor growth curve of the LNP-tRNA^Arg^_UGA_ group showed a significantly slower growth rate compared to the control groups after the first treatment (Day 8), resulting in a tumor volume reduction of approximately 70% and no mortality by the study endpoint (Day 22) (Figures 5C-5D and S17). In terms of functional p53 recovery, sup-tRNA treatment resulted in a 3.7-fold increase in full-length p53 expression and a 1.9-fold increase in downstream CDKN1A expression compared to vector treatment (Figures 5I and S18A). RT-qPCR analysis also revealed a 4.3-fold upregulation of *CDKN1A* transcripts in the sup-tRNA group relative to vector (Figures 5L and S18A). Consistently, LNP-tRNA^Gln^_UAG_ treatment also significantly attenuated tumor progression in the CRC Caco-2 xenograft harboring p53-E204*, with no treatment-related mortality observed by the endpoint (Day 50) (Figures 5E-5G and S17). Moreover, both protein levels of p53 and the mRNA levels of *CDKN1A* were markedly elevated following sup-tRNA treatment, whereas vector treatment showed minimal efficacy (Figures 5J-5M and S18B).

**Figure 5.**
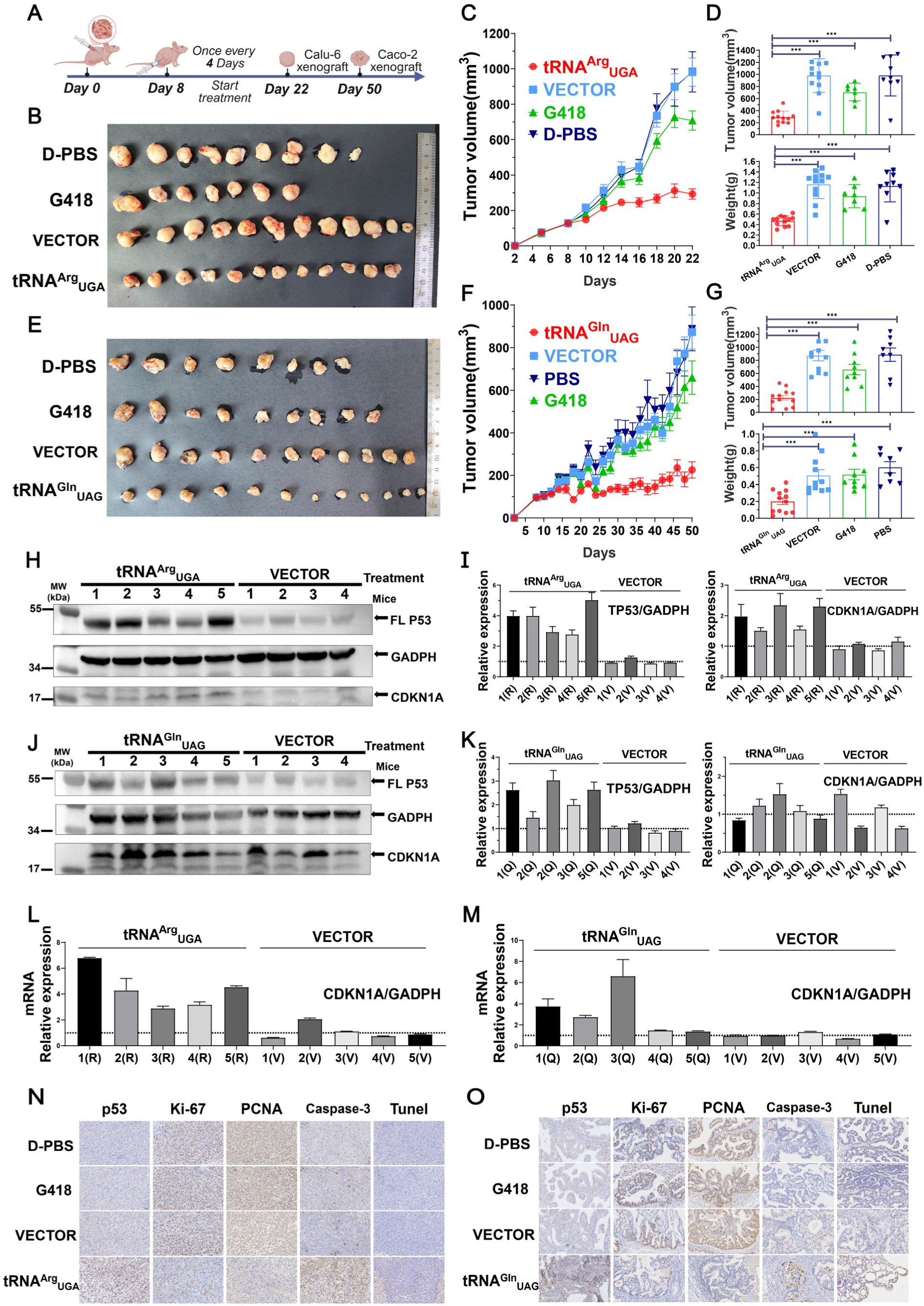
Therapeutic efficacy of sup-tRNA therapy *in vivo*. (A) Schematic of therapeutic evaluation of sup-tRNA therapy. (B) Images of Calu-6 xenograft excised tumors. (C and D) Tumor growth curves (C), volumes (above) and weights(below) of tumor collected on Day 22 (D) in Calu-6 xenograft model after tRNA^Arg^ treatment. *P*-values were calculated by one-sided t test. (E) Images of Caco-2 xenograft excised tumors. (F and G) Tumor growth curves (F), volumes (above) and weights(below) of tumor collected on Day 50 (G) in Caco-2 xenograft model after tRNA^Gln^_UAG_ treatment. *P*-values were calculated by one-sided t test. (H and I) Western blot analysis of full-length p53 recovery and protein expression of p53 target gene *CDKN1A* in Calu-6 xenograft treated with sup-tRNA (H) and quantification of the expression of each indicated protein relative to GAPDH expression, and normalized to the control mean (I). The abbreviation for tRNA^Arg^ is R, and the abbreviation for VECTOR is V. *P*-values were calculated by one-sided t test (J and K) Western blot analysis of full-length p53 recovery and protein expression of p53 target gene *CDKN1A* in Caco-2 xenograft treated with sup-tRNA (J) and quantification of the expression of each indicated protein relative to GAPDH expression, and normalized to the control mean (K). The abbreviation for tRNA^Gln^_UAG_ is Q, and the abbreviation for VECTOR is V. *P*-values were calculated by one-sided t test. (L and M) Quantification of mRNA levels of p53 target genes *CDKN1A* in Calu-6 (L) and Caco-2 (M) xenograft by RT-PCR. All the mRNA expression levels are normalized to *GAPDH*. Data are represented as mean ± SEM and individual data points shown for n = 3 independent biological replicates. *P*-values were calculated by one-sided t test. (N and O) Immunohistochemistry analysis of Calu-6 (O) and Caco-2 (P) xenograft including p53, proliferation and apoptosis markers after corresponding treatments. Scale bar=50 µm. Data shown here were representative examples from n=3 independent biological replicates.

The tumor regression induced by p53 restoration is primarily attributed to p53-mediated apoptosis and growth inhibition^32,33^, as evidenced by a reduction in Ki-67 and PCNA-positive proliferating cells, along with an increase in activated caspase-3 and TUNEL-positive apoptotic cells in both xenograft models following sup-tRNA treatment (Figures 5N-5O). Overall, these findings highlight the robust efficacy and excellent safety profile of p53 restoration through sup-tRNA-mediated translational readthrough, warranting further investigation into its broader application in cancer therapy.

### *In vivo* potential toxicity of sup-tRNA

Although aforementioned results showed that sup-tRNA alone did not inhibit cell growth in the NCI-H1299 (*TP53*-null) cell line, only when co-transfection of both sup-tRNA and *TP53* nonsense mutation gene led to growth inhibition (Figure 4B). However, further investigations are needed to determine whether the observed tumor regression is attributable to p53 restoration rather than potential sup-tRNA toxicity.

Next, the potential toxicity of sup-tRNA *in vivo* was assessed in a mouse xenograft model. Seven-week-old BALB/c mice were injected with an identical number of *TP53*-null 4T1 cells (Figure S21A). According to previous reports^19–22^, 4T1 and Calu-6 cells showed similar *in vivo* growth rates. Treatment of *TP53*-null xenografts with LNP-tRNA, tRNA^Arg^_UGA_ or tRNA^Gln^_UAG_, could not reduce tumor weight or volume by the study endpoint (Figures S21B-S21C), and the tumor growth curves showed no significant difference compared to the control group (Figure S21D). Moreover, no significant changes were observed in p53, Ki-67 and PCNA-positive proliferating cells, along with activated caspase-3 and TUNEL-positive apoptotic cells in each group (Figure S21F). Consistently, we could not find significant perturbations in the p53 signaling pathway after LNP-tRNA treatment (Figure S22). These results suggested that sup-tRNA could induce tumor regression by restoring tumor suppressor proteins, instead of its potential toxicity.

Furthermore, both in BALB/c nude mice and BALB/c mice, we did not observe any toxicity induced by LNP-tRNA. Safety assessments demonstrated no significant abnormalities in histopathological evidence of tissue damage (Figures S19 and S23A) or hematological parameters (Figures S20 and S23B) in sup-tRNA-treated mice, supporting the favorable therapeutic index of this approach.

## DISCUSSION

The distinct biological characteristics of oncogenes and tumor suppressor genes pose fundamentally different challenges for therapeutic intervention. The oncogene addiction in malignant tumors has a compelling therapeutic rationale for the development of targeted therapies^34,35^, such as kinase inhibitors^36^ and PROTACs^37^, which aim to inhibit or degrade hyperactive proteins. In contrast, tumor suppressor genes that often harbor loss-of-function mutations, typically require functional protein restoration, posing a significant pharmacological challenge^4,5^. Furthermore, mutant tumor suppressor proteins not only lose tumor-suppressive functions, but can also actively antagonize reintroduced wild-type proteins through dominant negative effect, limiting the therapeutic efficacy of gene therapy^11,12,38^. In this work, we showed that sup-tRNA could address these challenges by simultaneously restoring full-length p53 expression while converting dominant-negative truncations into functional variants. Notably, the generation of enhanced suppressor p53 mutants (Figures S14-S15) through specific amino acid substitution suggests a promising strategy to overcome the therapeutic limit imposed by the transcriptional activity of wild-type p53.

Moreover, we have developed a tailored framework for sup-tRNA in addressing ‘undruggable’ tumor suppressor mutations^4,5,10,39^. The computational pipeline, which incorporates analysis of readthrough efficiency and structural perturbation, could be adapted to other frequently mutated tumor suppressors such as RB1, PTEN, APC and STK-11 (Figures 1F-1G and S11). Unlike development of traditional small molecules targeting specific mutant proteins, the sup-tRNA strategy is not restricted by protein type and mutation type, as we presented that designed sup-tRNAs could overcome all three types of nonsense mutations across various tumor suppressor genes.

Here we report that the super-tRNA and superb-tRNA strategy could overcome nonsense mutations and restore neutral or advantageous p53 variants with enhanced anti-cancer function (Figure S15). An immense collection of genomic research has demonstrated that proteins can harbor rare advantageous mutations. However, neutral and advantageous mutations in *TP53* have received far less attention compared to extensively investigated deleterious mutations, despite it reports that approximately half of all p53 variants are neutral mutations^31^. This provides ample space for the application of the super-tRNA and superb-tRNA strategy. As the super-tRNA therapy for *TP53* nonsense mutations, we particularly recommend using tRNA^Gln^_UAG_ to suppress TAG nonsense codon where the original amino acid is glutamate or tryptophan, and using tRNA^Arg^_UGA_ to suppress TGA nonsense codon where the original amino acid is serine or tryptophan, with the assistance of computational tools. Significantly, we also remind that these strategies should be used with caution for the nonsense mutations at evolutionarily conserved residues.

Recent research have demonstrated that sup-tRNA exhibited extremely low or even no readthrough in normal termination codons, demonstrating the high safety profile of sup-tRNA^25,26^. Consistently, our study showed that sup-tRNA^Arg^_UGA_ and sup-tRNA^Gln^_UAG_ induced efficient readthrough in nonsense mutations with no detectable toxicity *in vitro* and *in vivo* (Figures S19-S23). However, in addition to the classical role of tRNAs as mRNA decoders, certain tRNAs could respond to external perturbations and regulate gene expression^40^ which either promotes or suppresses tumor progression^41,42^. As engineered from natural tRNAs, certain sup-tRNAs are in theory capable of driving or inhibiting cancer development beyond the readthrough of nonsense mutations^41,42^, suggesting excluding tumor proliferation-related tRNAs in sup-tRNA engineering for cancer therapy.

Overall, this study opens a new avenue for precision oncology that goes beyond the oncogene-centric paradigm. By translating the concept of tumor suppressor restoration into a viable therapeutic strategy, tRNA-based therapy could ultimately redefine treatment strategies for the majority of human cancers driven by suppressor protein inactivation.

## RESOURCE AVAILABILITY

### Lead contact

Requests for further information and resources should be directed to and will be fulfilled by the lead contact, Qing Xia.

### Materials availability

Requests for materials will be made available by the lead contact with a materials transfer agreement.

### Data and code availability

This paper does not report original code. Any additional information required to reanalyze the data reported in this paper is available from the lead contact upon request.

## ACKNOWLEDGMENTS

We are grateful to Y. Zhao and S. Shu for comments and feedbacks on this article. This work was financially supported by the National Key R&D Program of China, Frontier Biotechnology, (Grant No. 2023YFC3403300) and the National Natural Science Foundation of China (Grant No. U23A20106).

## AUTHOR CONTRIBUTIONS

Q.X. and B.K. conceived the idea and designed the experiments. B.K. and Z.Z. performed the experiments and offered the original data. B.K., Z.D., Y.Y., and Y.S. provided materials for plasmid cloning. B.K., Z.M., Y.L. and J.L. conducted the animal experiments. B.K., Z.Z. and Q.X. organized the results and wrote the manuscript. All authors participated in data discussion and interpretation. Q.X. supervised the study.

## DECLARATION OF INTERESTS

Q.X. and B.K. are inventors on patents regarding the suppressor tRNA for tumor suppressor protein recovery. Q.X. is the founder of QiXia Decode Biotechnology Co., Ltd. and holds equity in this company. The other authors declare no competing interests.

## STAR METHODS

### The tumor suppressor gene mutations

The tumor suppressor gene mutation data was downloaded from the COSMIC database (https://cancer.sanger.ac.uk/cosmic) and IntOGen database (https://www.intogen.org/). Data of amino acid types of nonsense mutations in tumor suppressor genes collected from the COSMIC database, including *TP53* (COSMIC gene: COSG501), *RB1* (COSMIC gene: COSG16), *PTEN* (COSMIC gene: COSG15) and *APC* (COSMIC gene: COSG257430).

### Plasmid construction

Tumor suppressor genes (*TP53, RB1, PTEN, APC*) were constructed in the pCAGGS-FLAG-GUSB-myc-T2A-GFP plasmid. pCAGGS vector has a unique promoter combination of chicken β-actin promoter and CMV enhancer (CAG promoter), enabling high-level expression in a wide range of cell types. Subsequently, targeted mutations were achieved by introducing primers containing mismatched bases during PCR to obtain plasmids with site-specific nonsense mutations. The tRNAs were constructed in pcDNA3.1(+) vectors and their expression was initiated by the 7SK promoter. tRNA gene sequences were compiled from GtRNA database (http://gtrnadb.ucsc.edu/). PG13-luc (wt p53 binding sites) plasmid (Addgene ID: 16442) were purchased from Addgene. We next changed the luciferase reporter to the mCherry reporter to obtain the plasmid PG13-mCherry (Addgene ID: 195291). The lactic acid probe pcDNA-FiLa-Cyt and pcDNA-FiLa-C-Cyt were gifts from Prof Yi Yang of East China University of Science and Technology.

### Luciferase assay

The dual luciferase reporter expresses the Firefly luciferase and Renilla luciferase (as transfection control), we inserted the TAG, TAA and TGA stop codons between the Firefly and Renilla reporter genes to obtain a dual-fluorescent reporter system with nonsense mutations. The nonsense mutation dual luciferase reporter plasmid was co-transfected with sup-tRNA plasmid into HEK293T cells. The luciferase was detected using the dual luciferase reporter assay kit (Promega, USA) after 48 hours of transfection.

### Cell culture and transfection

Human HEK293T cells (ATCC CRL-11268) were cultured in DMEM medium (Gibco) as suggested by the repository, supplemented with 10% fetal bovine serum (FBS), 100 U/mL penicillin, 100 mg/mL streptomycin.

Human Calu-6 cells (ATCC HTB-56) were cultured MEM medium (Gibco) as suggested by the repository, supplemented with 10% FBS, 100 U/mL penicillin, and 100 mg/mL streptomycin.

Human Caco-2 cells (ATCC HTB-37) were cultured MEM medium (Gibco) as suggested by the repository, supplemented with 20% FBS, 100 U/mL penicillin, and 100 mg/mL streptomycin.

4T1 cells (ATCC CRL-2539) and human NCI-H460 cells (ATCC HTB-177) were cultured RPMI 1640 medium (Gibco) as suggested by the repository, supplemented with 10% FBS, 100 U/mL penicillin, and 100 mg/mL streptomycin. Human A549 cells (ATCC CCL-185) were cultured Ham’s F-12K medium (Gibco) as suggested by the repository, supplemented with 10% FBS, 100 U/mL penicillin, and 100 mg/mL streptomycin. The concentration of G418 Sulfate used in the *in vitro* experiments was 100 ug/ml, which is supplied as a solution in water was obtained from Gibco (USA).

All cell lines were tested to be mycoplasma-free and validated by STR profiling. Plasmids were transfected using Lipo3000 and P3000 (Invitrogen). Cell transfections were performed with the supplier’s recommendations.

### Western blot analysis

The equivalent of 3×10^5^ cells was subjected to 10% SDS-PAGE before transferring the proteins to a 0.22um PVDF membrane through eBlot® L1 Fast Wet Transfer System (Genscript, USA). Membranes were incubated overnight at 4 °C in the presence of a 1/200 dilution of anti-p53 antibody (DO1) (Santa Cruz Biotechnology, Dallas, TX, USA; catalog number: sc-126), recognizing human p53. A 1/2000 dilution of anti-P21 antibody (Proteintech, China; catalog number: 10355-1-AP), a 1/100000 dilution of anti-GAPDH antibody (Proteintect, China; catalog number: 1E6D9). After three washes of the membrane in TBS-Tween, the membranes were exposed to a solution of horseradish peroxidase-conjugated secondary antibody for detection of mouse-raised or rabbit-raised antibody (Proteintect, China) at a dilution of 1/2000 (catalog number: SA00001-1, SA00001-2), respectively. Antibodies were then detected with BeyoECL Moon (P0018FS, Beyotime, China). Photographs were performed with 5200 (Tanon, Shanghai, China). The original unprocessed gel pictures are provided in the source data file. Western blot results were semi-quantitatively analyzed by ImageJ software. Western blot experiments were performed at least third in parallel and representative results were reported. The original western blot images are shown in Supplementary Figures.

### qRT-PCR analysis

1.5 ×10^5^ cells per well were seeded in 6-well plates in a final volume of 2 ml. On the following day, cells were transfected with different plasmids of the indicated drugs for 48 h. The RNA extraction was performed using the FastPure Cell/Tissue Total RNA Isolation Kit V2 (Vazyme, China) according to the manufacturer’s recommendations. RNA was quantified using a NanoDrop Spectrophotometer (Thermo Scientific, USA). cDNA was synthesized from RNA using Hifair® AdvanceFast One-step RT-gDNA Digestion SuperMix for qPCR (Yeasen, China). Real-time quantitative PCR was performed in the Hieff UNICON® Universal Blue qPCR SYBR Green Master Mix (Yeasen, China) using Roche LightCycler® 480 II (Roche, Switzerland) with the following primers:

*GAPDH* mRNA (sense: 5′- GGAGCGAGATCCCTCCAAAAT-3′; antisense: 5′- GGCTGTTGTCATACTTCTCATGG-3′).

*TP53* mRNA (sense: 5′- CAGCACATGACGGAGGTTGT-3′; antisense: 5′- TCATCCAAATACTCCACACGC -3′).

*CDKN1A* mRNA (sense: 5′- TGTCCGTCAGAACCCATGC -3′; antisense: 5′- AAAGTCGAAGTTCCATCGCTC -3′).

*BBC3* mRNA (sense: 5′- GCCAGATTTGTGAGACAAGAGG -3′; antisense: 5′- CAGGCACCTAATTGGGCTC -3′).

*GADD45A* mRNA (sense: 5′- GAGAGCAGAAGACCGAAAGGA -3′; antisense: 5′- CACAACACCACGTTATCGGG -3′).

*BAX* mRNA (sense: 5′- CCCGAGAGGTCTTTTTCCGAG -3′; antisense: 5′- CCAGCCCATGATGGTTCTGAT -3′).

*ZMAT3* mRNA (sense: 5′- AGAAGCCTTTTGGGCAGGAG -3′; antisense: 5′- TGCTGCATAGTAATTTCGGAGTT -3′).

*FAS* mRNA (sense: 5′- TCTGGTTCTTACGTCTGTTGC -3′; antisense: 5′- CTGTGCAGTCCCTAGCTTTCC -3′).

*PMAIP1* mRNA (sense: 5′- ACCAAGCCGGATTTGCGATT -3′; antisense: 5′- ACTTGCACTTGTTCCTCGTGG -3′).

The relative expression fold was calculated by the 2^−ΔΔCt^ equation.

### PCR array

Cell transfected, total RNA extracted and cDNA synthesized as described above. cDNA mixed with qPCR SYBR, added into the p53 PCR Array 96-well plate (WcGENE Biotech, WC-MRNA0117-H) to detect the expression of a set of p53-related genes.

### Colony formation assay

After 8 hours of transfection, we digested and resuspended the cells. We planted cancer cells into the six-well plate at a density of 1000 cells/well and then cultured them in a complete culture medium at 37°C for 7-10 days. After being gently washed in PBS twice, the cells were fixed with 4% paraformaldehyde fix solution for 30 min. Crystal Violet Staining Solution (Beyotime, China; catalog number: C0121) was applied for staining the fixed cells for 30 min. IncuCyte SX5 imaging system (Sartorius AG, Goettingen, Germany) was used to count the number of colonies, total colonies area, and individual clone area.

### Cell growth curve analysis

Cell transfected, digested, and resuspended as described above. For the cell counting, cells were inoculated into a 24-well plate with an appropriate number of cells, digested at 24h, 48h, 72h, and 96h after inoculation, and then counted by cell counter (Countstar BioTech, China). For the CCK-8 assays, cells were seeded into a 96-well plate at 2000 cells per well. According to the protocol of CCK-8 solution (Beyotime, China; catalog number: C0040), 10 µl of CCK-8 solution diluted in 100 µl of complete culture medium replaced the original medium of each group at indicated points. After the cells were incubated in the dark at 37 °C for an additional 1 h, we detected viable cells by using absorbance at a 450nm wavelength through Microplate Reader AMR-100 (Allsheng, China).

### The landscape of gene mutation

The STAR-counts data and mutation maf data of tumors were downloaded from the TCGA database (https://portal.gdc.cancer.gov). The data in TPM format was subsequently extracted from the STAR-counts data and normalized using log2(TPM+1). The samples that had RNAseq data, maf mutation data, and clinical information were then retained. The maftools package in R software was utilized to download and visualize the somatic mutations of patients. Statistical analysis was conducted using R software, version v4.0.3.

### Molecular dynamics simulation

The molecular dynamics simulations of all studied protein/DNA complexes were performed with the AMBER 22 molecular simulation package. The crystal structure of tumor-suppressor protein p53/DNA complex (PDB code: 2AC0) was used for the calculation. The starting structures of all molecular systems were solvated in TIP3P water using an octahedral box, which extended 8 Å away from any solute atom. To neutralize the negative charges of simulated molecules, Na+ counter-ions were placed next to each negative group. MD simulations were carried out by using the PMEMD module of AMBER 22. The calculation began with 500 steps of steepest descent followed by 500 steps of conjugate gradient minimization with a large constraint of 500 kcal mol-1 Å-2 on the atoms of the protein and DNA. Then 1000 steps of steepest descent followed by 4000 steps of conjugate gradient minimization with no restraint on the complex atoms were performed. Subsequently, after 200 ps of MD, during which the temperature was slowly raised from 0 to 300 K with weak (5 kcal mol-1Å-2) restraint on the protein/DNA, the final unrestrained production simulations of 200.0 ns were carried out at constant pressure (1 atm) and temperature (300 K). In the entire simulation, SHAKE was applied to all hydrogen atoms. Periodic boundary conditions with minimum image conventions were applied to calculate the non-bonded interactions. A cutoff of 10 Å was used for the Lennard–Jones interactions. The final respective conformations of all the protein/DNA complexes used for discussion were produced from the last 50.0 ns of MD.

### Whole-transcriptome analysis

Total RNA was extracted from tumor xenograft of different mice (three replicates for each group) using TRIzol reagent. RNA libraries were constructed using the VAHTS Universal V6 RNA-seq Library Prep Kit for Illumina (Vazyme, NR604-02) according to the manufacturer’s instructions. The concentration of the extracted RNA was detected using the Nanodrop2000 (Thermo Fisher, Nanodrop2000), and its integrity was tested using the Agilent 2100 (Vazyme, G2939BA). The quality and quantity of the constructed libraries were assessed using the BioAnalyzer 2100 system (Agilent Technologies).

Subsequently, the libraries were sequenced on an Illumina NovaSeq X Plus platform with 150-bp paired-end reads. The sequencing data were quality-controlled using the Q30 metric. After adapter trimming at the 3’ end and removing low-quality reads with the fastp(https://github.com/OpenGene/fastp, fastp-V0.20.1), high-quality clean reads were aligned to the reference genome using the HISAT2(https://daehwankimlab.github.io/hisat2/, hisat2-2.0.4).

Guided by the gffcompare, the ballgown software package was used to obtain gene-level FPKM (Fragments Per Kilobase of transcript per Million mapped reads) values as the expression profiles of mRNA. The total number of expressed genes and the log2FPKM values of mRNA in different tumor xenograft groups were plotted and compared. Significant differences between samples were analyzed using DEseq2, and genes with a multiplicity of difference FoldChange>2 times and P-value<0.01 were defined as differential genes and subsequently analyzed.

### Preparation and characterization of LNP-tRNAs

The ionizable lipid SM-102 was purchased from Guanshan Nano-Tech (China). The 1,2-distearoyl-sn-glycero-3-phosphocholine (DSPC) was purchased from Avanti Polar Lipids (USA); 1,2-Dimyristoyl-rac-glycero-3-methylpolyoxyethy lene-2000 (DMG-PEG2000) was purchased from NOF Corporation (Japan); and cholesterol (Chol) was purchased from Sigma-Aldrich (USA).

The molar ratio of SM102 LNPs was SM-102: DSPC: Cholesterol: DMG-PEG=50:10:38.5:1.5. Nanoassemblr (Precision Nanosystems, Canada) and Hamilton liquid handling system (Microlab STAR, USA) were used to prepare LNPs at three volumetric portions of tRNA to 1 volumetric portion of lipid solutions to obtain a lipid concentration of 2.81 mg/mL. The clean-up of the crude formulations was achieved by dialysis in PBS using Slide-A-Lyzer G2 dialysis cassettes (Thermo Scientifc, USA).

Transmission Electron Microscope (TEM) image of LNPs was obtained by HT7800 (HITACHI, Japan). The hydrodynamic size and zeta potential of LNPs were measured by Zetasizer (Malvern, USA).

### Xenograft tumor model

Animal experiments were proven by the Institutional Animal Care and Use Committee of Peking University Health Science Center (Animal Protocol ID: DLASBD0468). Female BALB/c nude mice or BALB/c mice were housed in an SPF facility in a 12-h light-dark cycle and allowed free access to water and food. At 7 weeks of age, BALB/c nude mice were subcutaneously inoculated with 5×10^6^ calu-6 cells or 1×10^7^ caco-2 cells into the left flank, both cell lines in 50% BD Matrigel™. BALB/c mice were subcutaneously inoculated with 1×10^6^ 4T1 cells into the left flank.

Tumor volumes and mice weight were calculated every 2 days, tumor volumes using the following standard formula: volume = 0.5 × length × width^2^. When the tumor volume reached about 200 mm^3^, treatment was initiated using either LNP-tRNA (10ug tRNA/injection), LNP VECTOR, G418 (20 mg/kg/ injection) or D-PBS. Treatments were made by intratumor injection once every 4 days. When the tumor volume reaches 800mm^3^, the animal is considered dead and *Kapplan-Mayer* survival curves were plotted for survival analysis. Maximum allowed weight loss was 10% or maximum tumor volume 1000 mm^3^. Mice were sacrificed after treatment. Tumors were dissected, weighed, and cut in thirds: one-third was frozen in liquid nitrogen for protein analysis, one-third was kept in the RNAlater (Beyotime, China; catalog number: R0118) for RT-qPCR analysis and one-third was fixed in 4% paraformaldehyde (PFA) for histological analysis and immunostaining.

### Immunofluorescence staining

Collected tumors were fixed in 4% PFA at room temperature and prepared as 10-μm tumor sections. For p53 and CDKN1A, antigens were repaired by Tris-EDTA antigen repair solution G1206 (Servicebio, Wuhan, China). For ki67, PCNA and caspase-3, the antigen was repaired by Citric Acid Antigen Repair Solution G1202 (Servicebio, Wuhan, China). For quenching of endogenous peroxidase, the sections were incubated in 3% hydrogen peroxide solution for 25 min at room temperature, protected from light, and the slides were placed in PBS (pH 7.4) and washed three times for 5 min each time by shaking on a shaker. 3% BSA was then added to cover the tissue evenly and closed for 30 min at room temperature. The blocking solution was removed and sections were incubated with diluted primary antibody at 4°C overnight in a wet box, 1/50 dilution of anti-p53 antibody (DO1) (Santa Cruz Biotechnology, Dallas, TX, USA; catalog number: sc-126), recognizing human p53. 1/200 dilution of anti-P21 antibody (Proteintect, China; catalog number: 10355-1-AP). 1/200 dilution of anti-ki67 antibody (Servicebio, China; catalog number: GB121141). 1/200 dilution of anti-PCNA antibody (Servicebio, China; catalog number: GB12010). A 1/200 dilution of anti-cas3 antibody (Servicebio, China; catalog number: GB11532). After washed in PBS on a decolorizing shaker with shaking three times, 5 min each time, sections were shaken dry and the tissue was covered with a drop of secondary antibody (HRP-labelled) of the corresponding species to the primary antibody and incubated for 50 min at room temperature. The slides were subsequently shaken dry after washing and freshly prepared DAB color development solution G1212 (Servicebio, Wuhan, China) was added dropwise, while the time of color development was controlled under the microscope. The sections were counterstained in hematoxylin followed by dehydration with graded ethanol, n-butanol, and xylene, then the sections were taken out of xylene to dry and seal. The slides were photographed under pathology slide scanner ws-10 (Wisleap, China). NDPView2 software (Hamamatsu, Japan) was used to analyze and record these images.

### Histological analysis

Blood samples from mice were collected from the orbital vein, and stored in the tube with K2-ethylenediaminetetraacetic acid (EDTA) at 4°C. The hematological parameters were analyzed using a BC-2800Vet® Auto Hematology Analyzer (Shenzhen Mindray Bio-Medical Electronics Co., Ltd, Hamburg, Germany). The heart, liver, spleen, lung, and kidney tissues from different groups of mice were isolated and fixed in 4% PFA. They were then sequentially dehydrated in 70%, 95%, and 100% ethanol, followed by defatting with xylene for 2 hours before being embedded in paraffin. The 10-μm-thick sections were obtained and subjected to H&E staining. The slices were examined using a pathology slide scanner (WS-10, Wisleap, China). Image analysis and recording were performed using NDPView2 software (Hamamatsu, Japan).

### Statistical analysis

Data were analyzed using the GraphPad Prism v5 statistical software package (GraphPad Software, La Jolla, CA). Data were expressed as means ± SD or SEM. The one-way analysis of variance (ANOVA) with Newman-Keuls multiple comparisons test with statistical significance set at P < 0.001 was performed. Appropriate statistical tests have been mentioned in Figure legends.

## Notes

### Summary of Updates

All figures revised and supplemental files updated to improve readability. Section on Methods to clarify lipid nanoparticles delivery of sup-tRNA.

