## Supplemental figures for "Restoration of tumor suppressor protein with enhanced activity using super-tRNAs to induce tumor regression"

A

| Dependent variables | (1) Cancer types where is driver | (2) Cohorts where is driver | (3) Cancer types where is driver | (4) Cohorts where is driver | (5) Cancer types where is driver |
|---|---|---|---|---|---|
| *Nonsense_COSMIC* | **0.586*****<br>**(5.36)** | **1.451*****<br>**(5.12)** | | | |
| *Missense_COSMIC* | | | **0.039**<br>**(1.11)** | **0.097**<br>**(0.93)** | |
| *Synonymous_COSMIC* | | | | | **-0.222*****<br>**(-2.68)** |
| Constant | 0.881*<br>(1.68) | -1.191<br>(-0.93) | 2.924**<br>(2.21) | 3.840<br>(1.00) | 7.189***<br>(5.96) |
| Observations | 314 | 314 | 314 | 314 | 314 |
| R-squared | 0.250 | 0.228 | 0.005 | 0.005 | 0.036 |

Robust t-statistics in parentheses
*** p<0.01, ** p<0.05, * p<0.1

| Dependent variables | (1) Cancer types where is driver | (2) Cohorts where is driver | (3) Cancer types where is driver | (4) Cohorts where is driver | (5) Cancer types where is driver |
|---|---|---|---|---|---|
| *Truncating _IntOGen* | **0.237*****<br>**(5.47)** | **0.614*****<br>**(5.38)** | | | |
| *Missense_IntOGen* | | | **-0.236*****<br>**(-4.13)** | **-0.570*****<br>**(-3.74)** | |
| *Synonymous_IntOGen* | | | | | **-0.375*****<br>**(-4.34)** |
| Constant | 0.643<br>(1.14) | -2.241<br>(-1.63) | 18.923***<br>(5.25) | 42.622***<br>(4.49) | 12.206***<br>(6.07) |
| Observations | 314 | 314 | 314 | 314 | 314 |
| R-squared | 0.187 | 0.187 | 0.104 | 0.091 | 0.147 |

Robust t-statistics in parentheses
*** p<0.01, ** p<0.05, * p<0.1

B

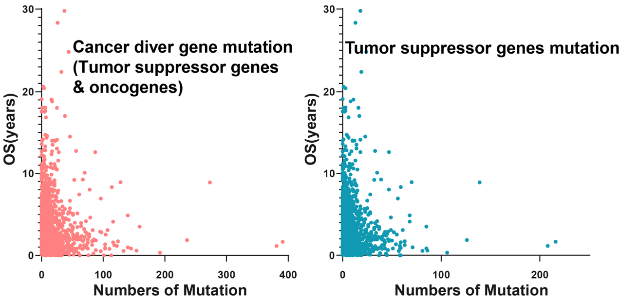


C

| Variables | (1) *Time_all* | (2) *Time_lof* |
|---|---|---|
| *Num* | **-0.007***<br>**(-1.786)** | **-0.027*****<br>**(-3.328)** |
| Constant | 1.956***<br>(37.694) | 1.967***<br>(50.539) |
| Observations | 2,412 | 2,407 |
| R | 0.001 | 0.005 |

**Figure S1. Identification and analysis of driver tumor suppressor gene mutations**

(A) OLS analysis between the proportion of mutation types of tumor suppressor genes and their driver ability. Robust t-statistics are in parentheses. The rate of tumor suppressor gene nonsense/truncation mutations data is derived from IntOGen & COSMIC, and the number of driver cancer types and cohorts of tumor suppressor gene data is derived from IntOGen.

(B and C) Correlation analysis between driver mutation burden and survival. Scatter plots (B) and Ordinary Least Square (OLS) analysis (C) show negative correlations between total driver mutations (x-axis) and overall survival (OS, y-axis, years), with emphasis on tumor suppressor gene (TSG) mutations. Survival data from TCGA cohort (n=10,641).

A

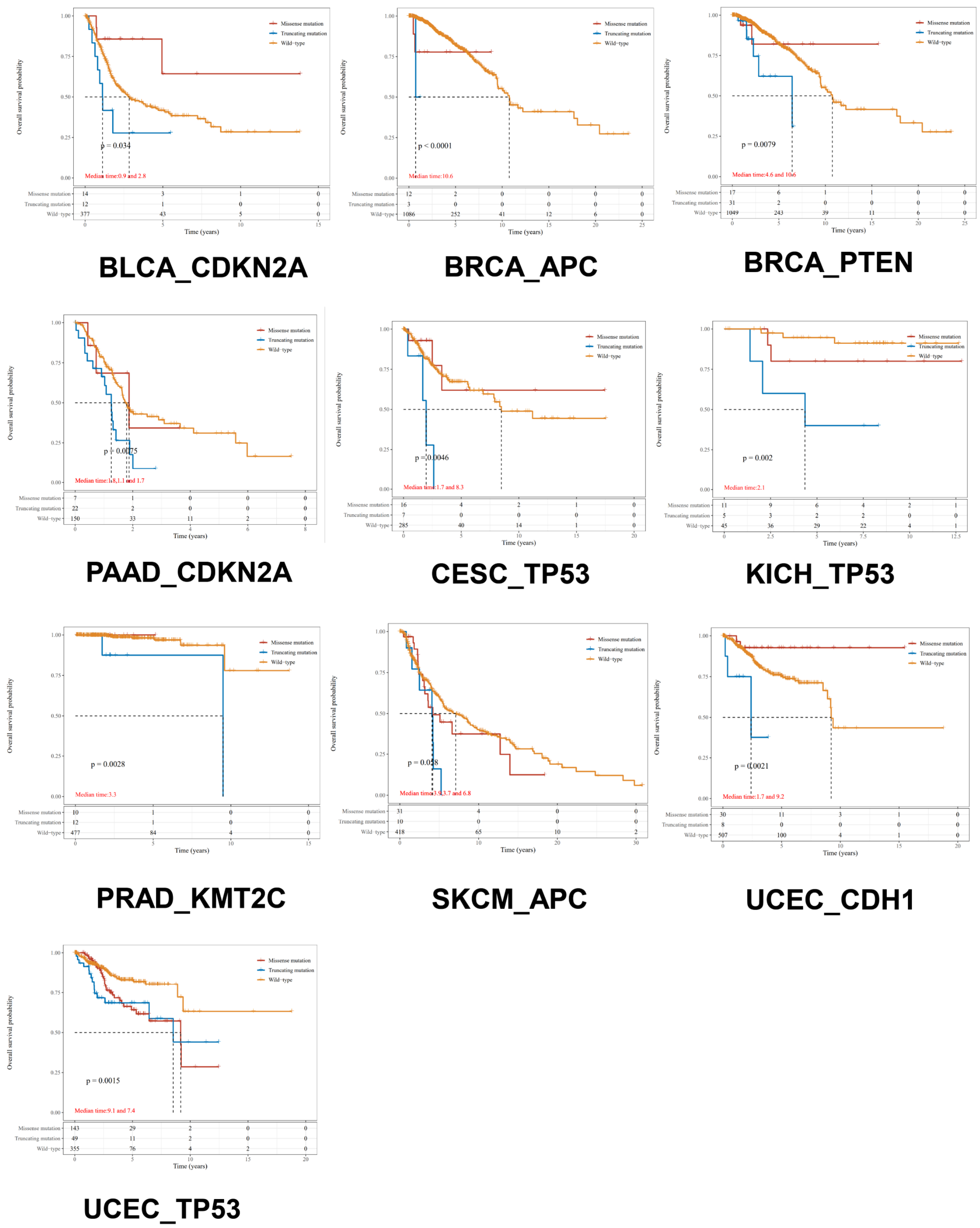


**Figure S2. Truncating mutations of tumor suppressor genes in specific cancer types lead to poorer prognosis**

(A) The KM survival curve distribution of different tumor suppressor gene mutation types groups in the TCGA dataset, with log-rank tests conducted between groups. The 95% CI represents the confidence interval for the HR. The median time represents the time corresponding to a 50% survival rate for different groups (i.e. median survival time), measured in years. The numbers in the table below represent the number of patients still in the study at a given time. The cancer types and tumor suppressor genes are labeled below the picture.

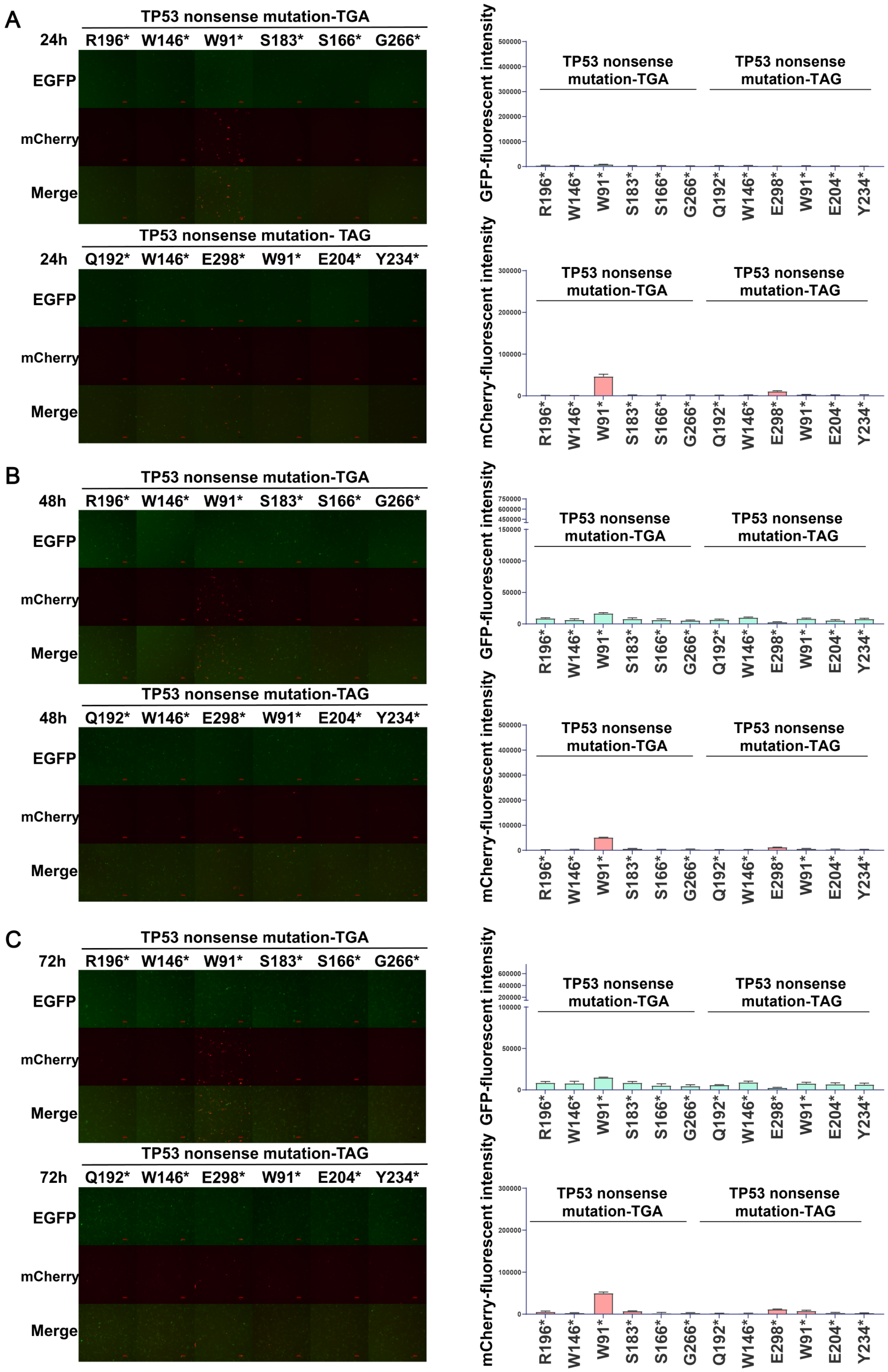


**Figure S3. TP53 nonsense mutations lead to the loss of p53 protein expression and function**

(A, B and C) Representative microscopy images (left) and quantification of GFP and mCherry positive cells (right) show p53 expression and function in NCI-H1299 cells with 12 kinds of nonsense mutations after 24h(A), 48h(B) and 72h(C) transfection. Data are mean ± s.e.m of three biological replicates. Scale bars = 100 μm.

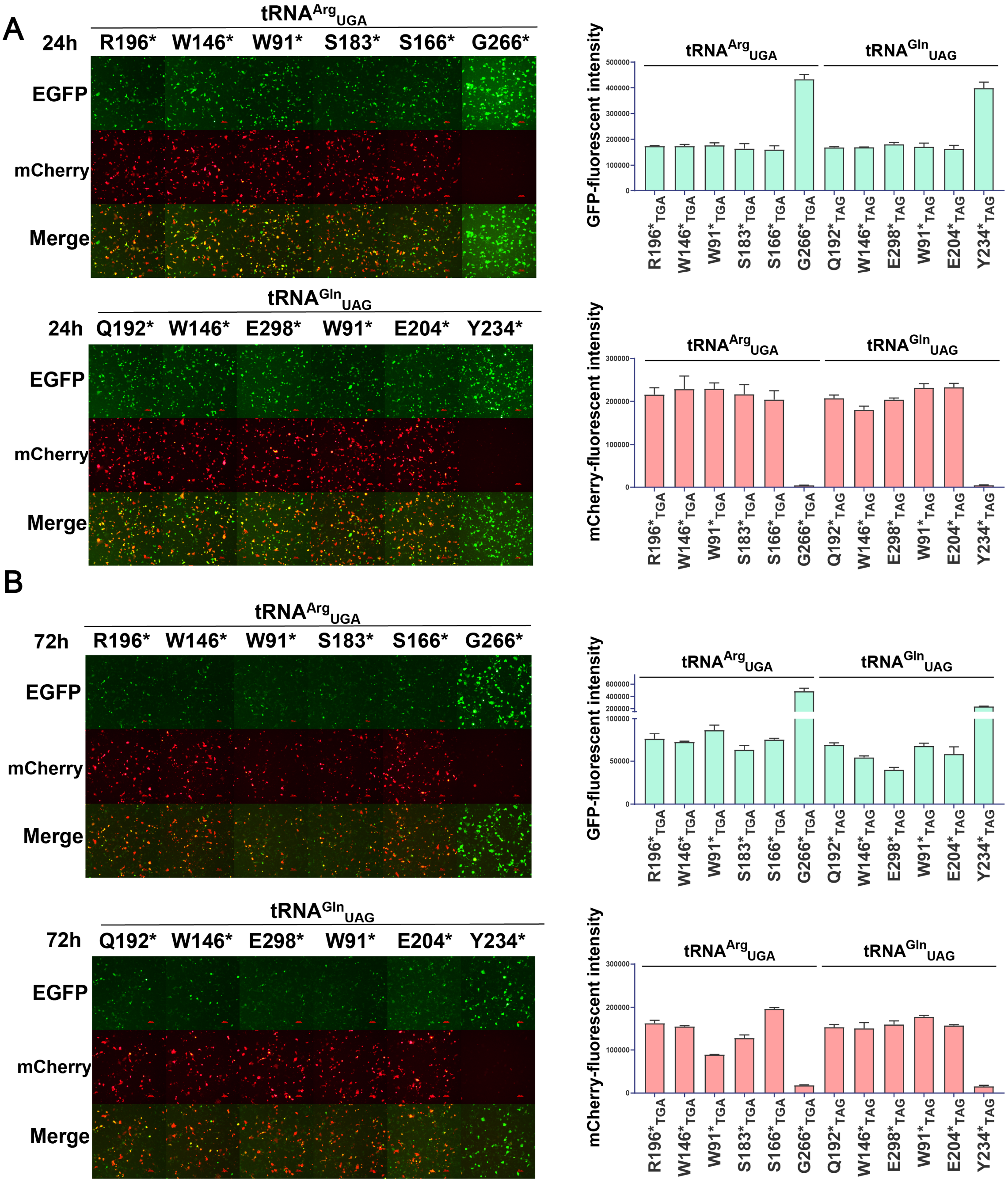


**Figure S4. tRNA$^{Arg}$$_{UGA}$ and tRNA$^{Gln}$$_{UAG}$ restore the expression and function of p53 protein from different nonsense mutations**

(A and B) Representative microscopy images (left) and quantification of GFP and mCherry positive cells (right) show p53 expression and function in NCI-H1299 cells with 12 kinds of nonsense mutations after tRNA $^{Arg}$$_{UGA}$ and tRNA$^{Gln}$$_{UAG}$ readthrough after 24h(A) and 72h(B) transfection. Data are mean ± s.e.m of three biological replicates. Scale bars = 100 μm.

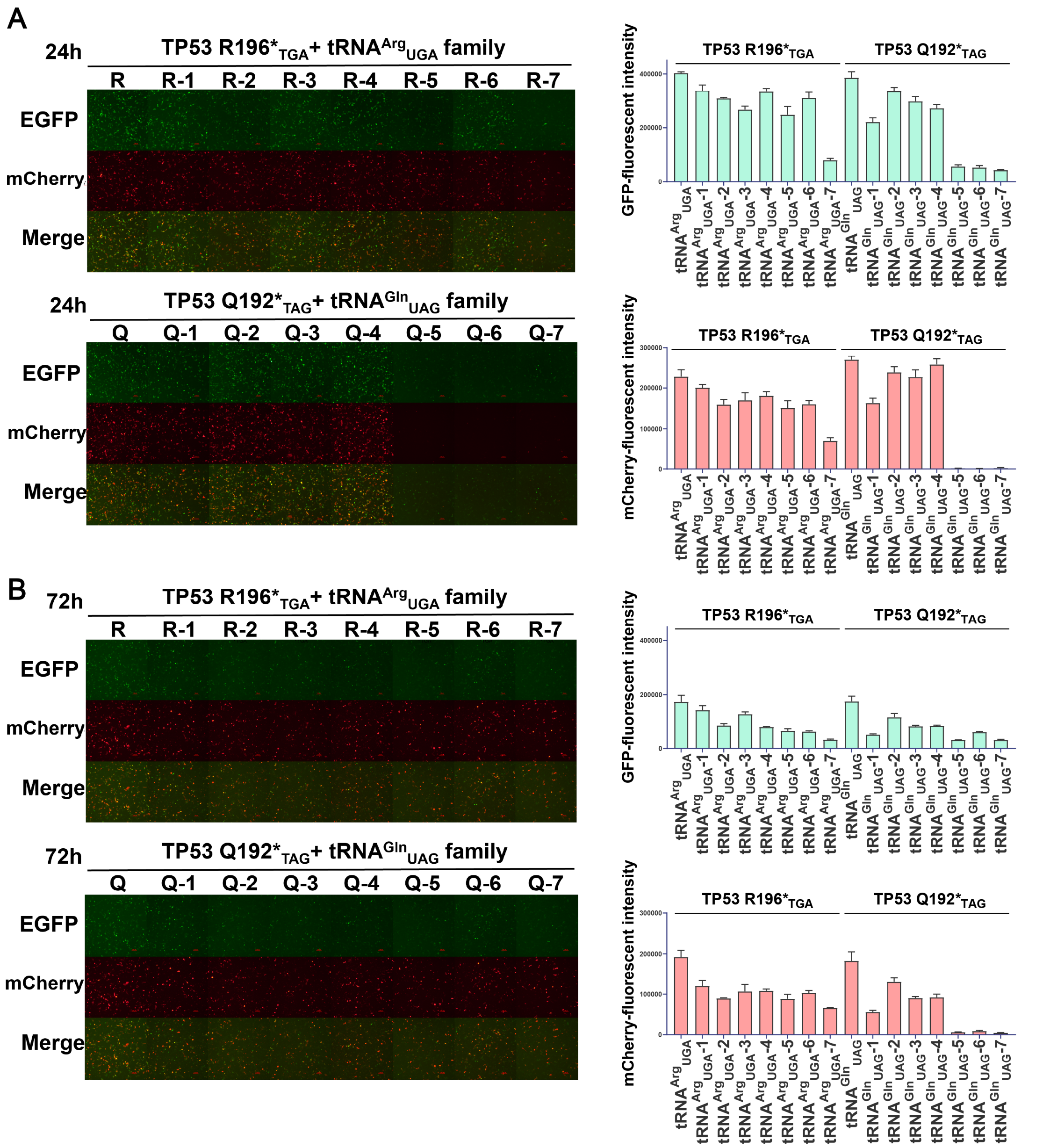


**Figure S5. tRNA$^{Arg}_{UGA}$ family and tRNA$^{Gln}_{UAG}$ family restore the expression and function of p53 protein from *TP53* R196*$_{TGA}$ and Q192*$_{TAG}$**

(A and B) Representative microscopy images (left) and quantification of GFP and mCherry positive cells (right) show p53 expression and function in NCI-H1299 cells with *TP53* R196*& *TP53* Q192* nonsense mutations after tRNA$^{Arg}_{UGA}$ family and tRNA$^{Gln}_{UAG}$ family sup-tRNAs readthrough after 24h(A) and 72h(B) transfection. The abbreviation of Arg, Gln is R, Q. Data are mean ± s.e.m of three biological replicates. Scale bars = 100 μm.

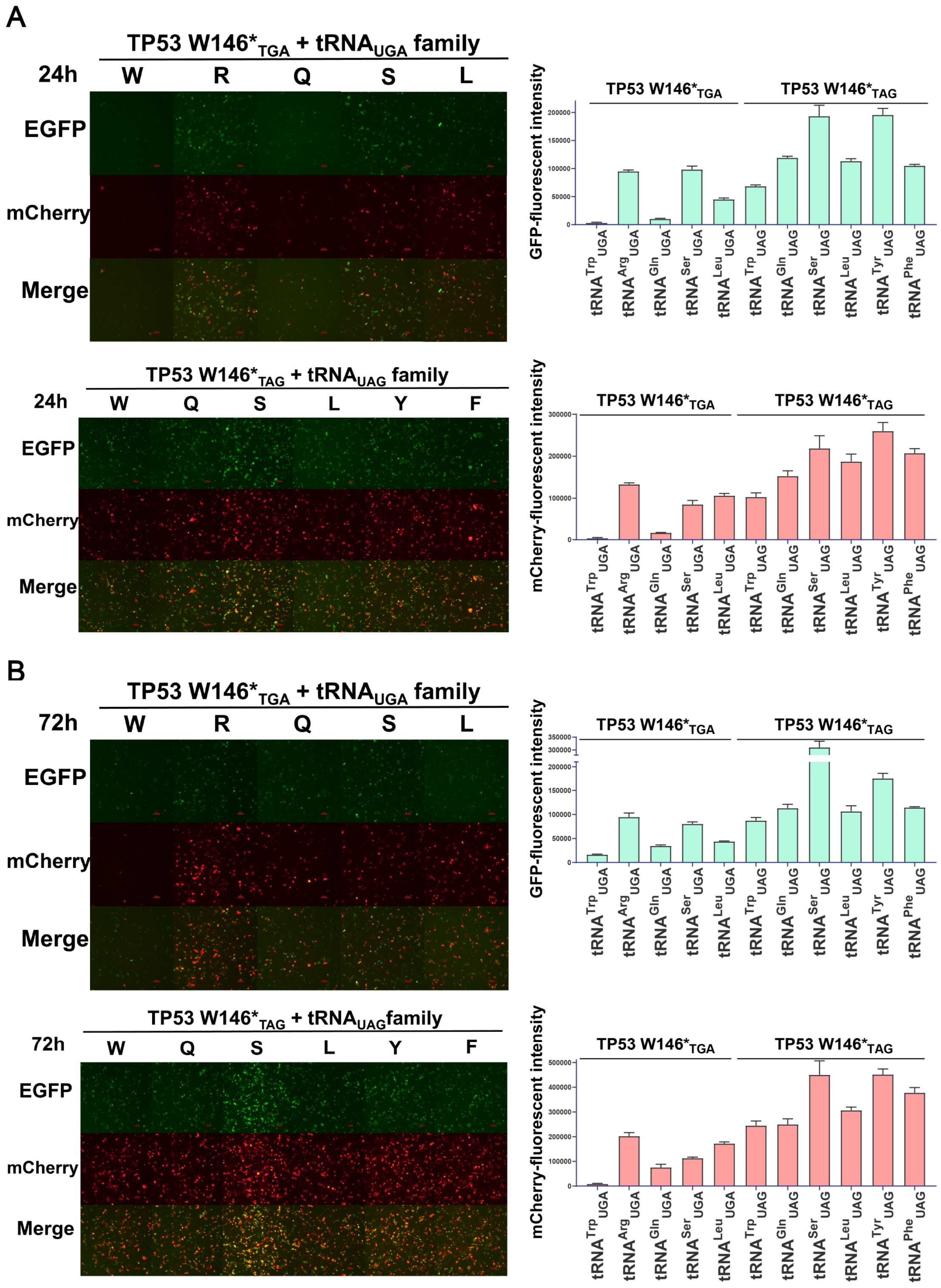


**Figure S6. tRNA$_{UGA}$ family and tRNA$_{UAG}$ family restore the expression and function of p53 protein from *TP53* W146*$_{TGA}$ and W146*$_{TAG}$**

(A and B) Representative microscopy images (left) and quantification of GFP and mCherry positive cells (right) show p53 expression and function in NCI-H1299 cells with *TP53* W146* nonsense mutations of tRNA$_{UGA}$ family and tRNA$_{UAG}$ readthrough after 24h(A) and 72h(B) transfection. The abbreviation of Trp, Arg, Gln, Ser, Leu, Tyr, Phe is W, R, Q, S, L, Y, F. Data are mean ± s.e.m of three biological replicates. Scale bars = 100 μm.

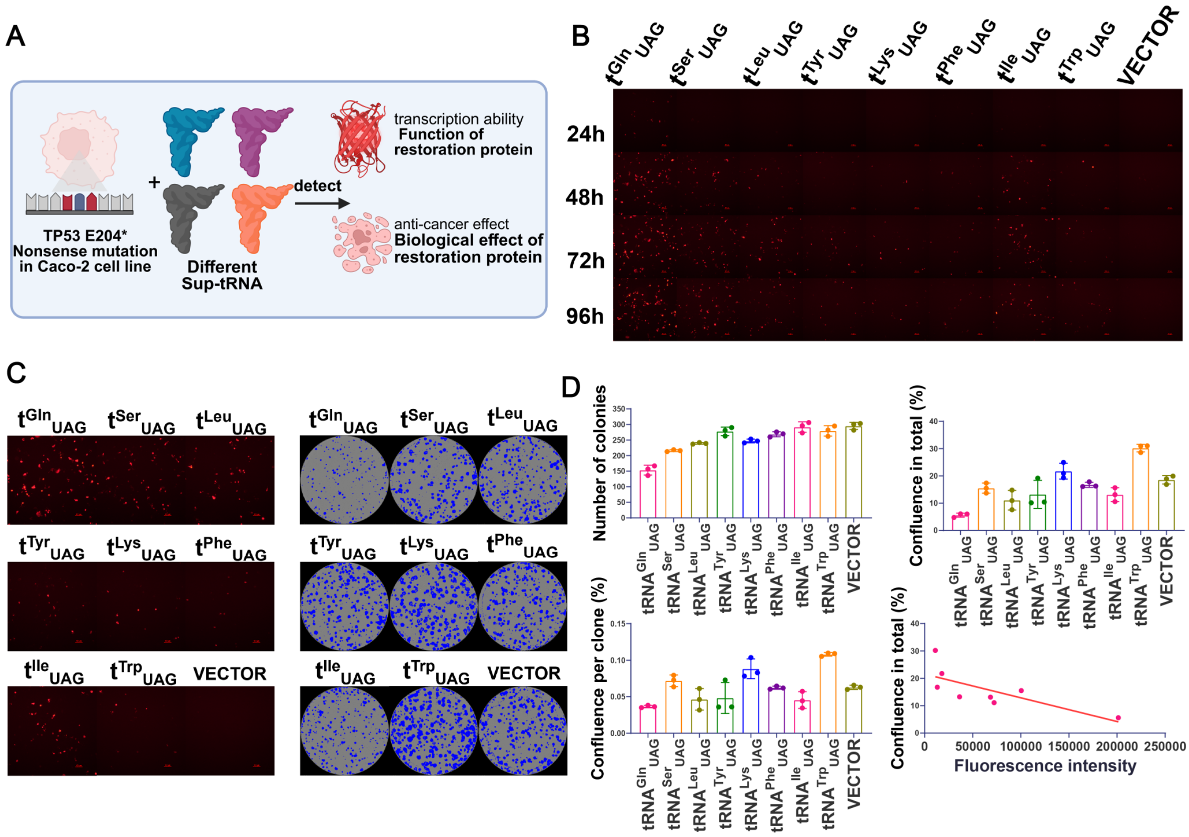


**Figure S7. Different sup-tRNAs elicits various degrees of antiproliferation activity through various degrees of p53 function restoration**

(A) Schematic of a genetically encoded reporter & colony formation system to measure the function and antiproliferation activity of restored p53 via different sup-tRNAs readthrough.

(B) Representative fluorescence images show various degrees of transcription function recovery of endogenous p53 in Caco-2 cells after 8 kinds of high-efficiency sup-tRNA readthrough at different time points. The abbreviation of tRNA is t. Scale bar=100 μm. Data shown here were representative examples from n=3 independent biological replicates.

(C and D) The 72h representative fluorescence images and corresponding colony formation image(C) & data(D) after 8 kinds of high-efficiency sup-tRNAs readthrough. Colonies are stained with crystal violet staining solution and counted after 10-12 days of culture. Scale bar=100 μm. Colony images are acquired by Incucyte SX5 and processed and analyzed by Incucyte 2022B Rev2. Data are represented as mean ± SEM, n = 3. The abbreviation of tRNA is t.

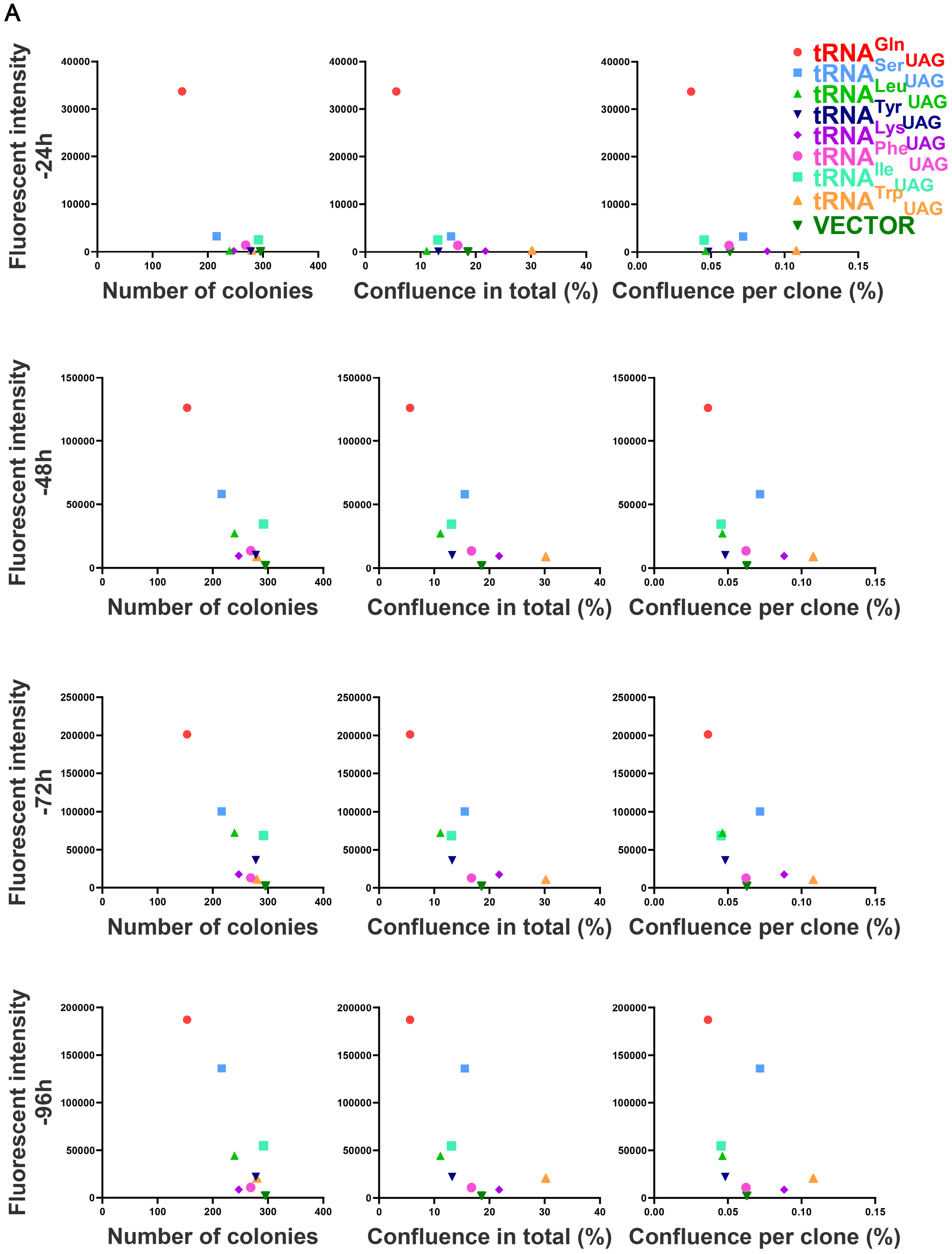


**Figure S8. Sup-tRNA-induced antiproliferation activity and the degrees of p53 function restoration are negatively correlated**

(A) Scatter plots of the quantitative data of fluorescent and colony formation at different time points after different sup-tRNA treatments.

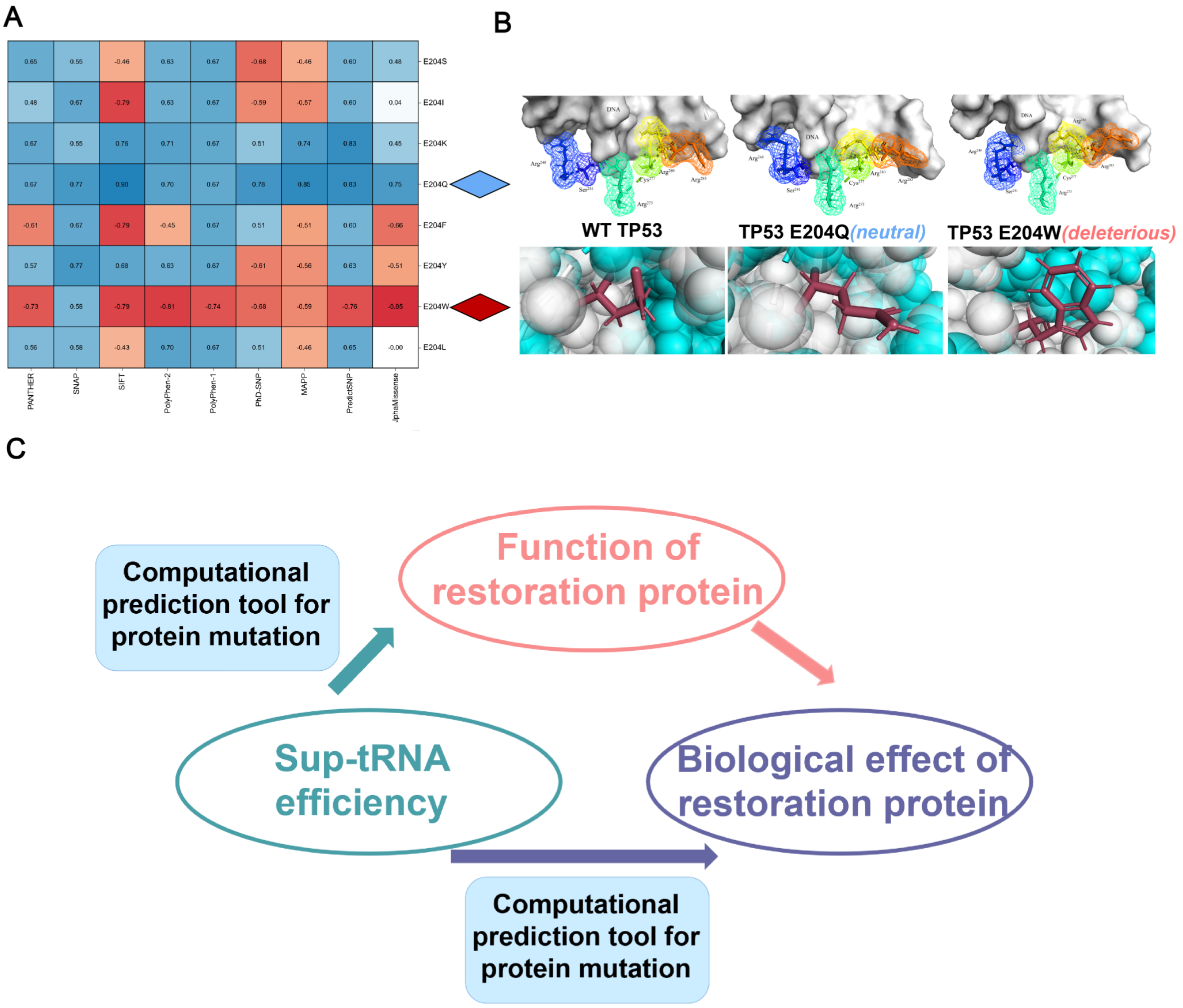


**Figure S9. Predictions of neutral/deleterious mutations assisted by computational tools and the correlation of the three elements of tRNA therapy**

(A) Predictive and functional profile of p53 E204 variants. The heat map corresponds to the predictive impact of each p53 variant ranging from -1 (red) to 1 (blue), with the lowest score being the most deleterious. Scores from the 9 algorithms in the panels including AlphaMissense, MAPP, PANTHER, PhD-SNP, PolyPhen-1, PolyPhen-2, SIFT, and SNAP. Red arrows indicate previous verified deleterious p53 variant E204W. Blue arrows indicate previous verified neutral p53 variant E204Q.

(B) Verified deleterious and neutral p53 variants mutant site amino acid residues. The above panels show the 204Q, and 204W structures of wild-type (WT) p53 DBD (2AC0). The panels below show the DNA binding peptides and DNA close-up view of the wild-type p53, neutral variant p53 E204Q and deleterious variant p53 E204W. The side chains of the six classic DNA binding residues are shown as sticks.

(C) Summary chart about establishment of sup-tRNA readthrough efficiency, function of restoration protein and biological effect of restoration protein three associations.

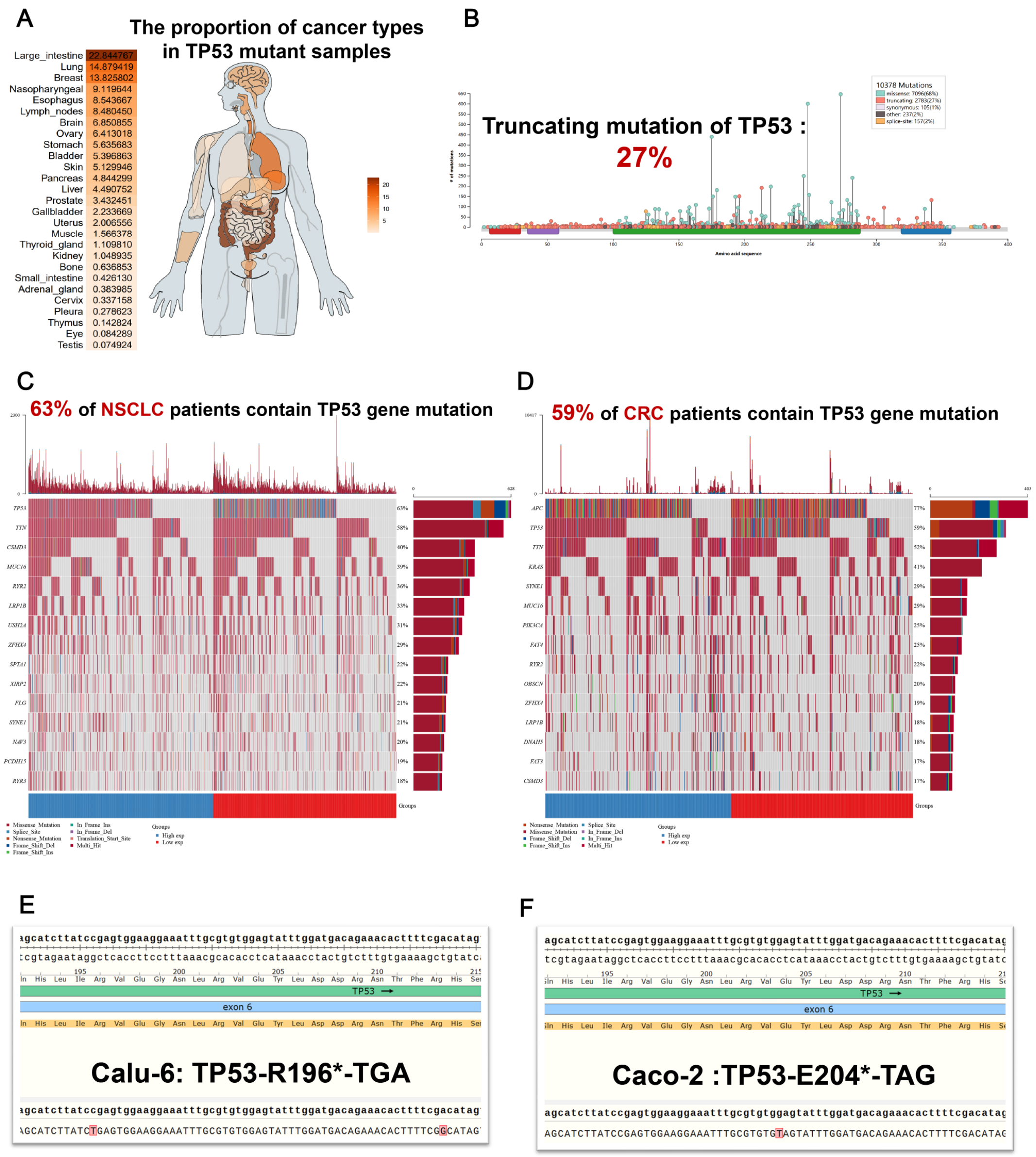


**Figure S10. TP53 mutation analysis and cell lines validation**

(A) Proportion of cancer species in *TP53* mutations samples. Total *TP53* mutation dates are downloaded from the COSMIC database.

(B) Lollipop diagram showing the mutation distribution of the *TP53* exon gene, where the mutation type and rate are indicated by the right of the figure. The TP53 mutation data were extracted from IntOGen while truncating mutations were marked in red.

(C and D) Oncoplot showing the somatic mutation landscape of the NSCLC (C) and CRC (D) tumor cohort. Genes are sorted by mutation frequency and samples are sorted by disease histology, as indicated in the annotation bar at the bottom. The sidebar graph shows the types and numbers of gene mutations. The waterfall graph shows the mutation information for each gene in each sample, with different colors with specific annotations at the bottom representing different mutation types.

(E and F) Sequencing verified the presence of genomic TP53 nonsense mutations in Calu-6 (E) and Caco-2 (F) cell lines.

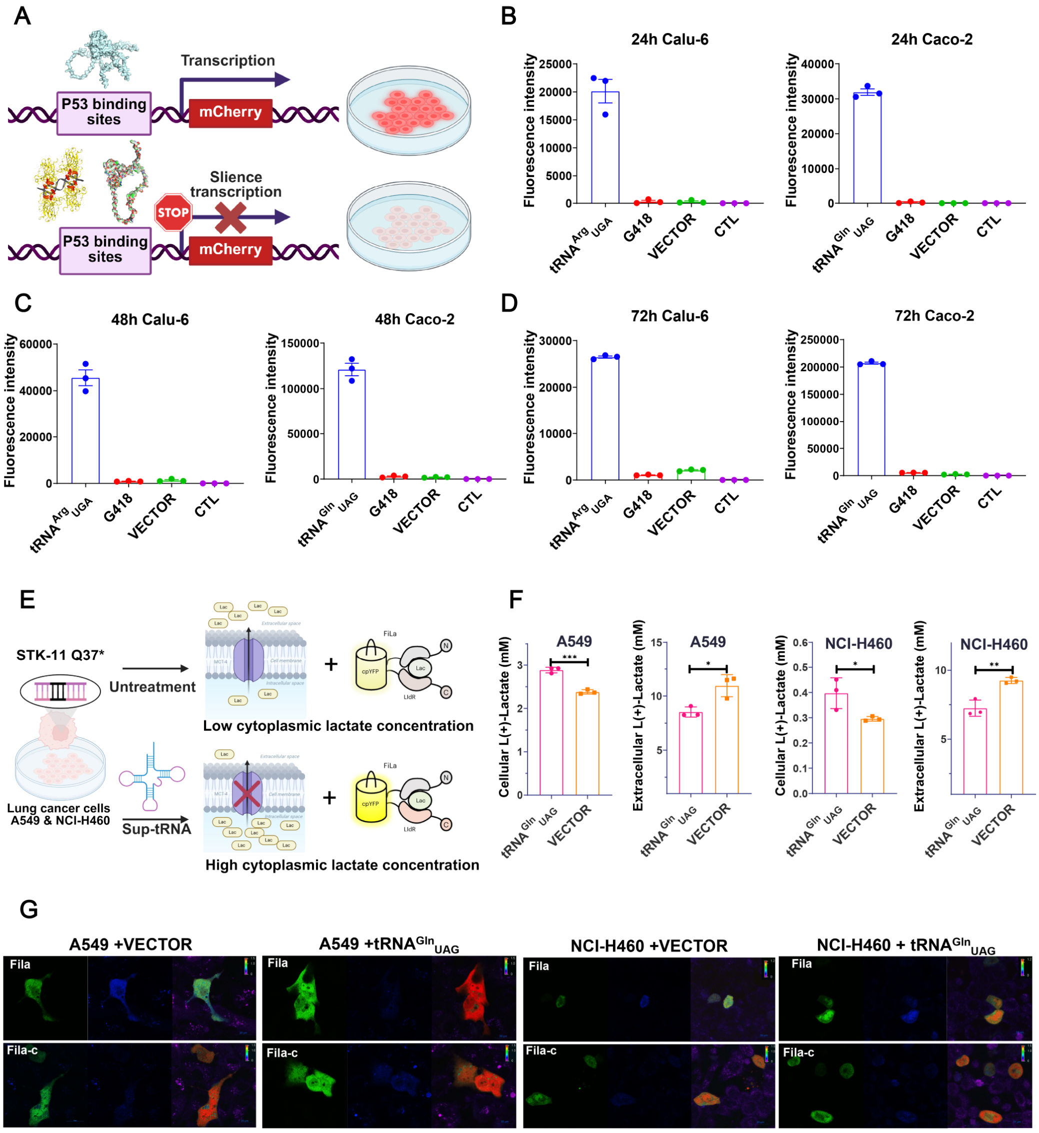


**Figure S11. Genetically encoded fluorescent sensing probes report restoration of p53 and STK-11 proteins from nonsense mutations after sup-tRNA treatment**

(A) Workflow about verification of functional p53 recovery by genetically encoded sensor (PG13-mCherry). (B, C and D) Quantification of fluorescence images shows the transcription function recovery of endogenous p53 in Calu-6 & Caco-2 cells after 24h(B), 48h(C) and 72h(D) sup-tRNA transfection. Data are mean ± s.e.m of three biological replicates.

(E) Workflow about verification of functional STK-11 recovery by genetically encoded lactate sensor (Fila).

(F) L (+)-lactate content in A549 & NCI-H460 (both *STK-11* Q36* cells) cytoplasm and medium was measured by Lactate content assay kit after $tRNA^{Gln}{}_{UAG}$ treatment. Data are mean ± s.e.m of three biological replicates

(G) Fluorescence images of FiLa in A549 & NCI-H460 after $tRNA^{Gln}{}_{UAG}$ treatment. Images were pseudo-colored by R488/405. Scale bar=20 μm. Data shown here were representative examples from n=3 independent biological replicates.

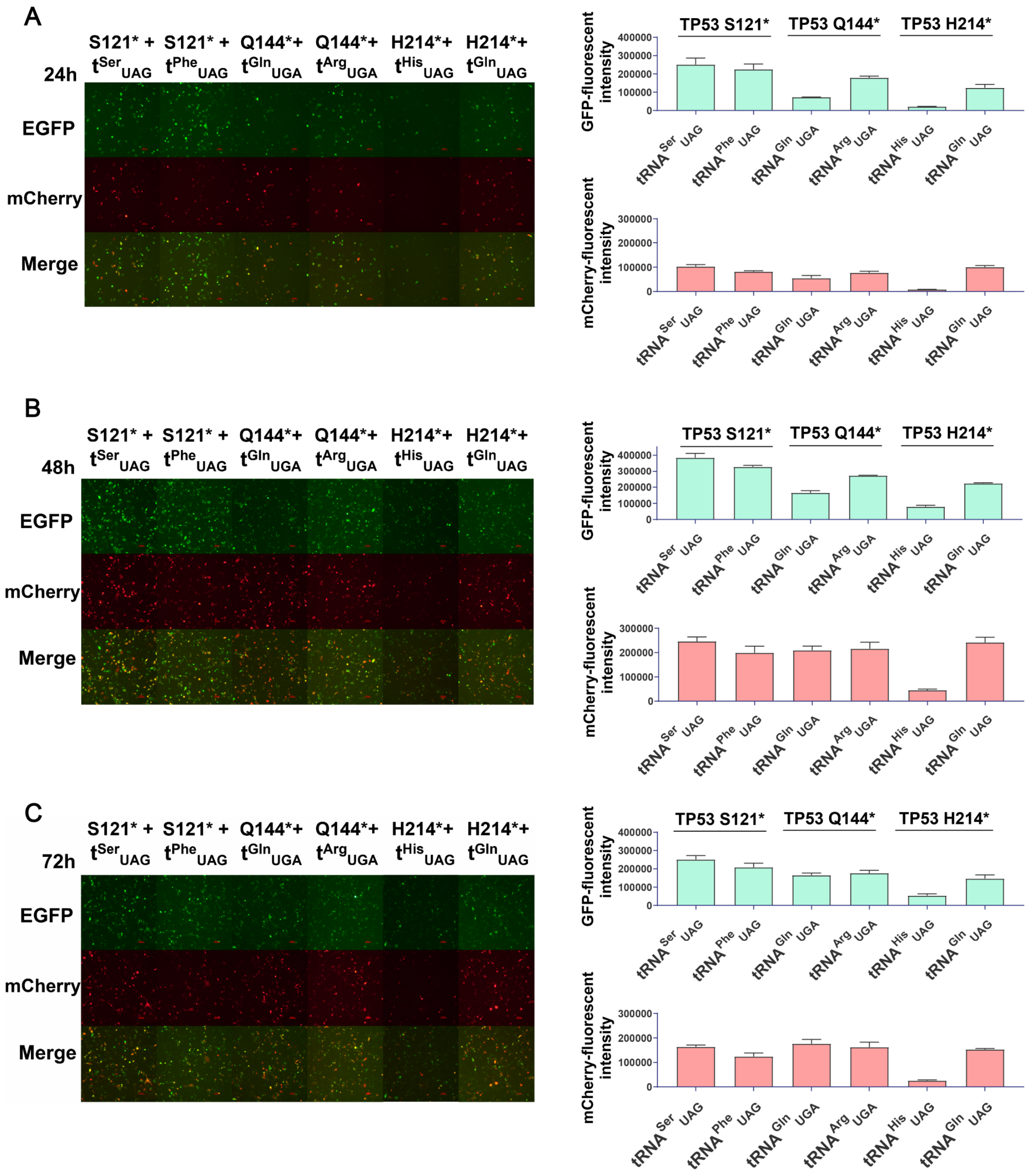


**Figure S12. The superb-tRNA restores the expression and function of p53 protein from TP53 S121*, Q144* and H214***

(A, B and C) Representative microscopy images (left) and quantification of GFP and mCherry positive cells (right) show p53 expression and function in NCI-H1299 cells with *TP53* nonsense mutations (S121*, Q144*, H214*) after specific sup-tRNAs readthrough after 24h(A), 48h(B) and 72h(C) transfection. The abbreviation of tRNA is t. Data are mean ± s.e.m of three biological replicates. Scale bars = 100 μm.

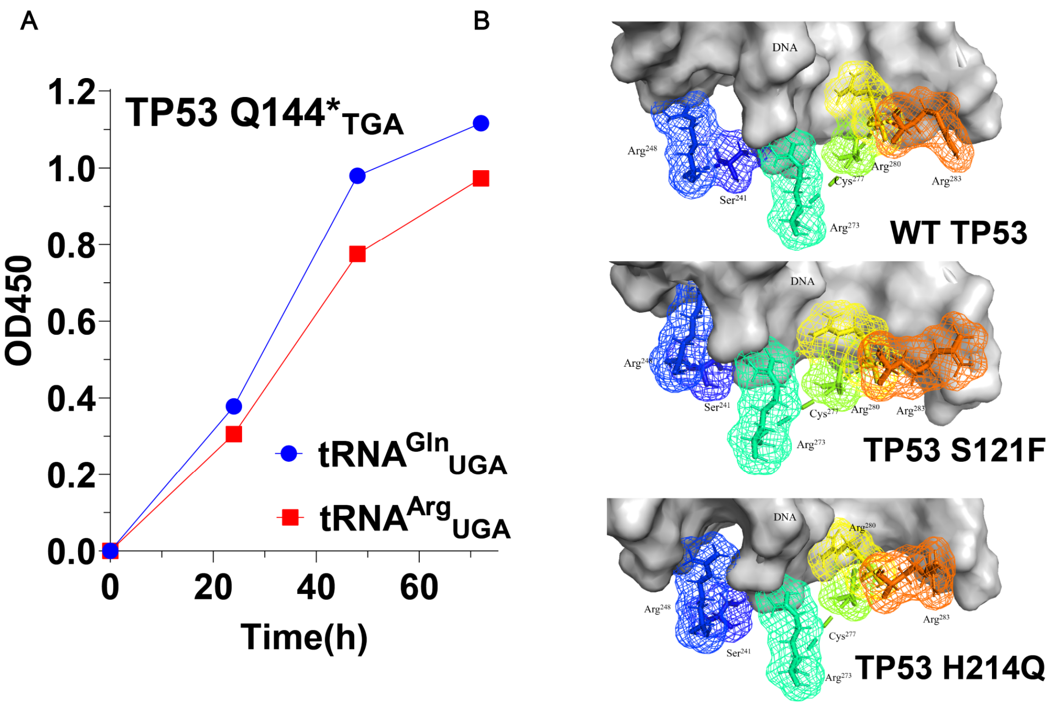


**Figure S13. Cell proliferation curve and structure simulation of the superb-tRNA therapy**

(A) Cell proliferation curve of the superb-tRNA therapy was measured by CCK-8. After transfecting cell for 8 hours, the NCI-H1299 cells are digested, spread in plates and further measured. Data are represented as mean, n = 3.

(B) The panels below close-up view of the p53 DNA binding peptides and DNA. including wild-type p53, enhance p53 variants S121F & H214Q. The side chains of the six classic DNA binding residues are shown as sticks.

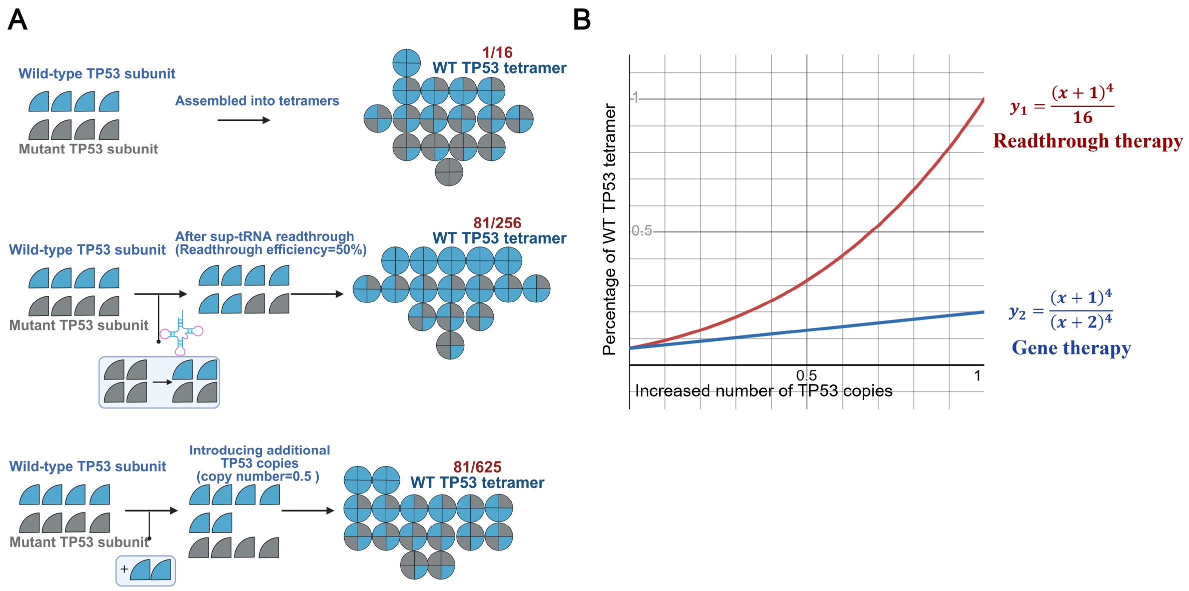


**Figure S14. tRNA therapy can better overcome the dominant negative (DN) effect compared to gene therapy**

(A and B) Workflow(A) and mathematical fit(B) about tRNA therapy can convert mutant p53 subunit into wt p53 subunit, thereby forming more wt p53 tetramers than gene therapy.

A

| | A. Restore to the original amino acids? | B. Protein expression | C. Protein function | D. Biological effect |
|---|---|---|---|---|
| Super-tRNA | No | Higher | Original | Higher |
| Super-tRNA (Superb-tRNA) | No | Higher | Higher | Higher |
| Super-tRNA (Superb-tRNA) | No | Original | Higher | Higher |
| Primary-sup-tRNA | Yes | Original | Original | Original |
| Lower-sup-tRNA | No | Lower | Lower | Lower |

eg: When the states in columns B and C are inconsistent, the judgment will base on column D.

B

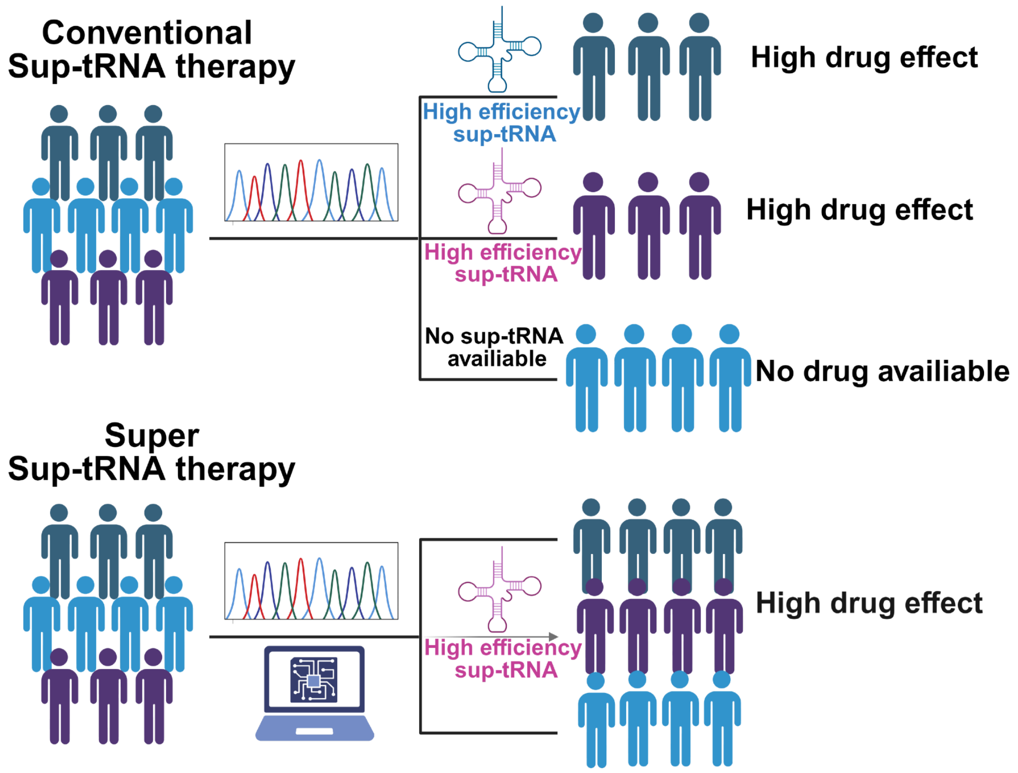


**Figure S15. The definitions of super-tRNA and superb-tRNA therapy**

(A) Summary chart about the classification and definitions of super-tRNA, superb-tRNA, primary-sup-tRNA and lower-tRNA therapy

(B) Compared with conventional sup-tRNA therapy, super sup-tRNA therapy can expand the patient population for the indications and enhance the therapeutic effect.

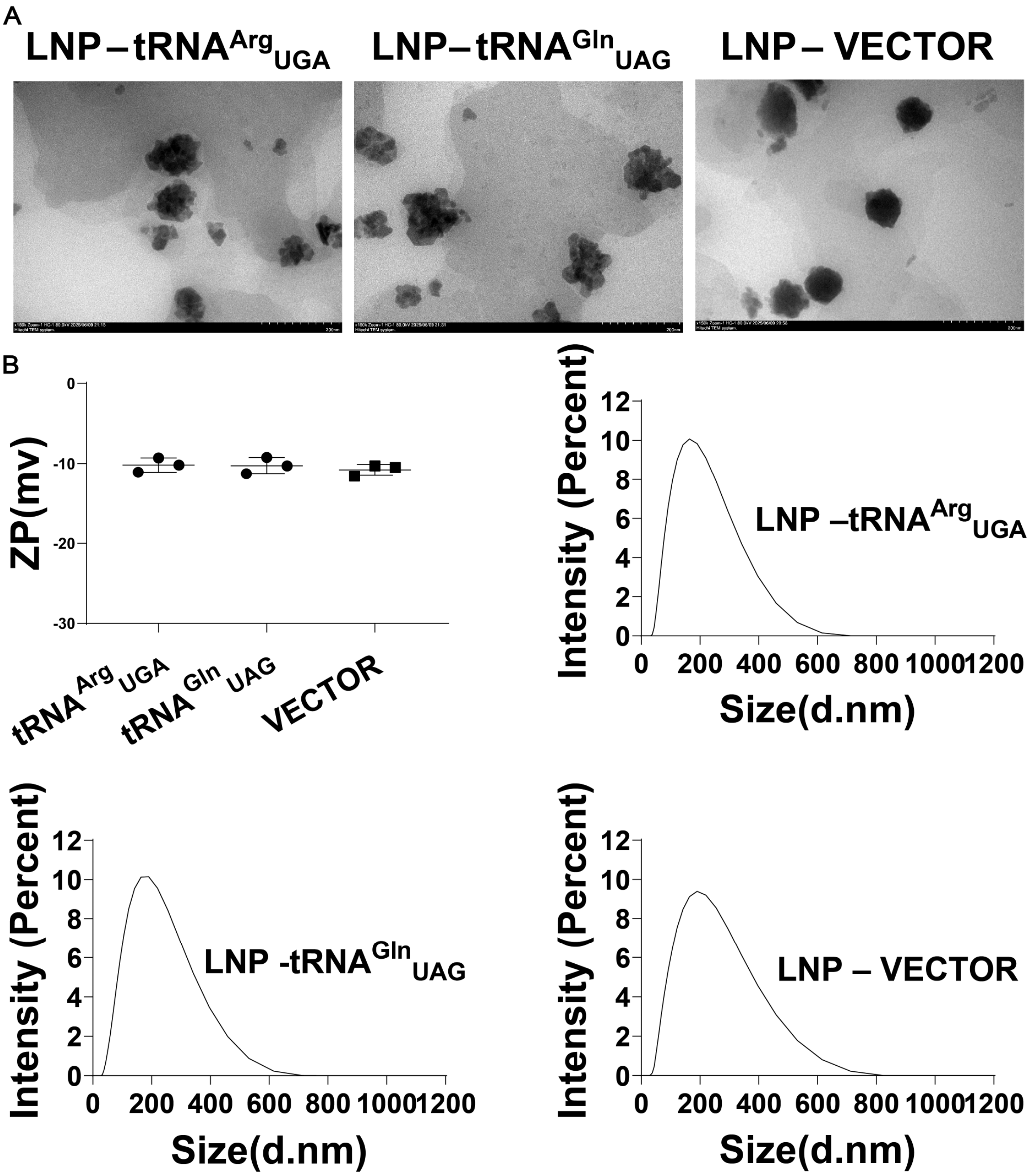


**Figure S16. Characterization of LNP-tRNAs**

(A) TEM images of LNP-tRNA$^{Arg}_{UGA}$, LNP-tRNA$^{Gln}_{UAG}$ and LNP-VECTOR. Scale bars = 200 nm

(B) Analysis of the zeta potential (top left corner) and size distribution and of LNP-tRNA$^{Arg}_{UGA}$, LNP-tRNA$^{Gln}_{UAG}$ and LNP-VECTOR. Data are mean ± s.e.m of three biological replicates.

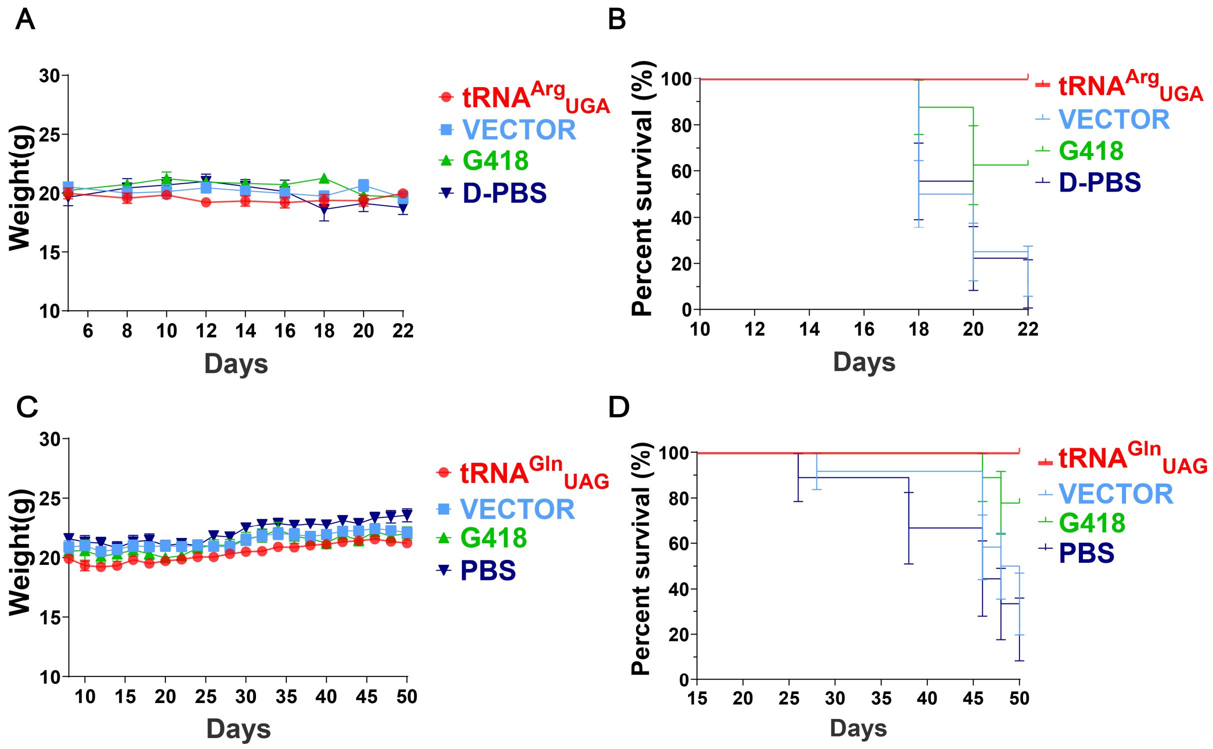


**Figure S17. Weight and survival curves of BALB/c nude after sup-tRNA treatment**

(A and B) Weight curve (A), and Kaplan-Meier survival curves (B) in Calu-6 xenograft model after LNP-$tRNA^{Arg}_{UGA}$ treatment.

(C and D) Weight curve (C), and Kaplan-Meier survival curves (D) in Caco-2 xenograft model after LNP-$tRNA^{Gln}_{UAG}$ treatment.

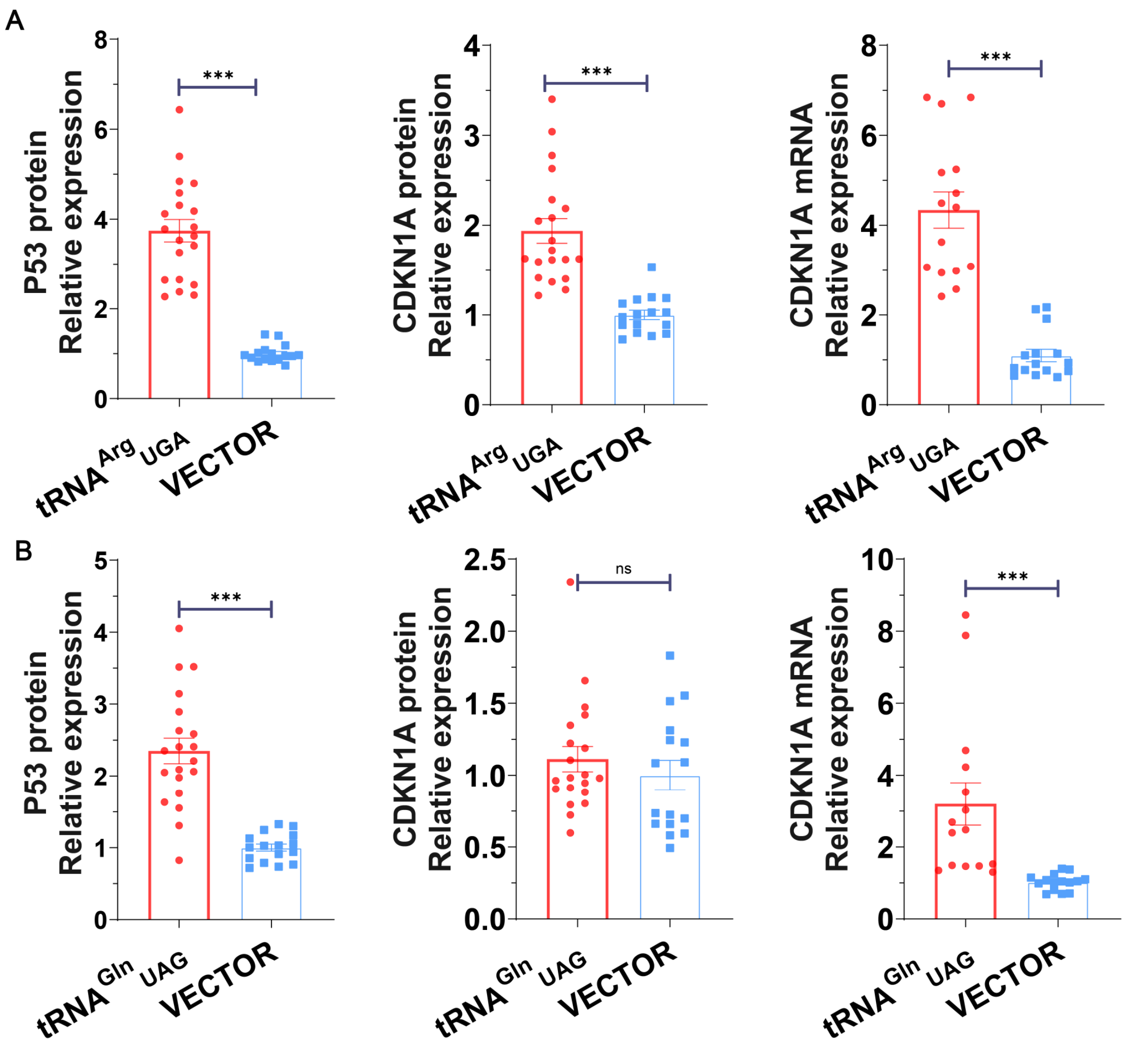


**Figure S18. Analysis of protein and mRNA expression in sup-tRNA in *vivo* treatment**

(A) Analysis of p53 (left) & CDKN1A protein (middle) and CDKN1A mRNA (right) expression in Calu-6 xenograft treated with tRNA$^{Arg}_{UGA}$. P-values were calculated by one-sided t test.

(B) Analysis of p53 (left) & CDKN1A protein (middle) and CDKN1A mRNA (right) expression in Caco-2 xenograft treated with tRNA$^{Gln}_{UAG}$. P-values were calculated by one-sided t test.

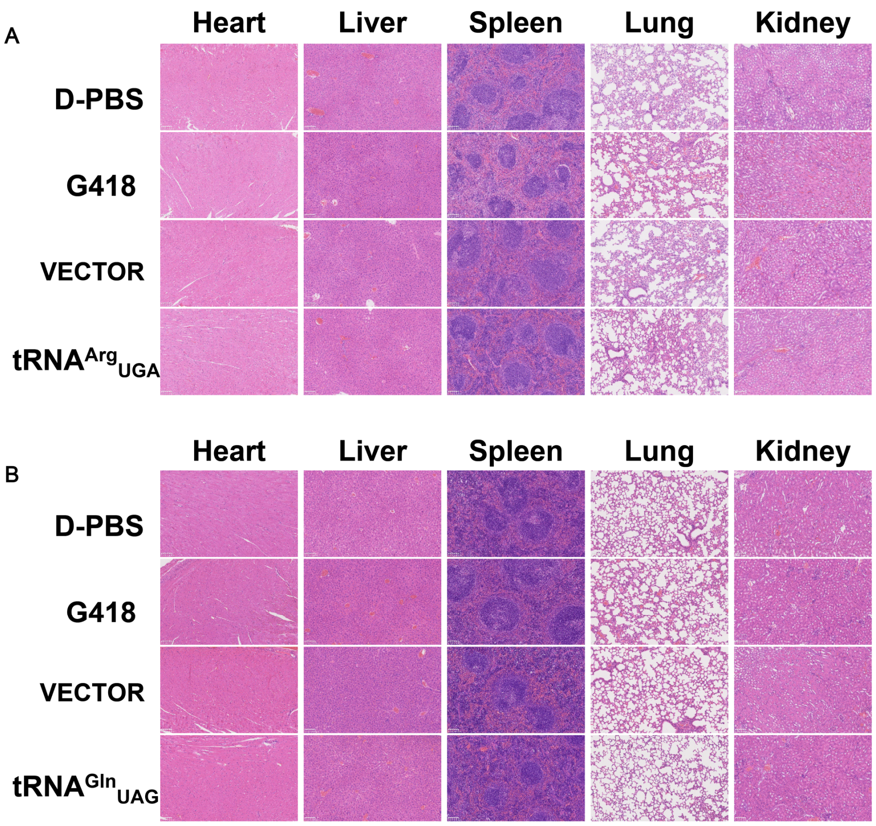


**Figure S19. H&E staining of the organs in sup-tRNA in-*vivo* treatment**

(A and B) Safety of the sup-tRNA treatment on heart, liver, spleed, lung, and kidney of Calu-6 (A) and Caco-2 (B) xenograft nude mice assessed by H&E staining. Scale bars = 100 μm. Data shown here were representative examples from n=3 independent biological replicates.

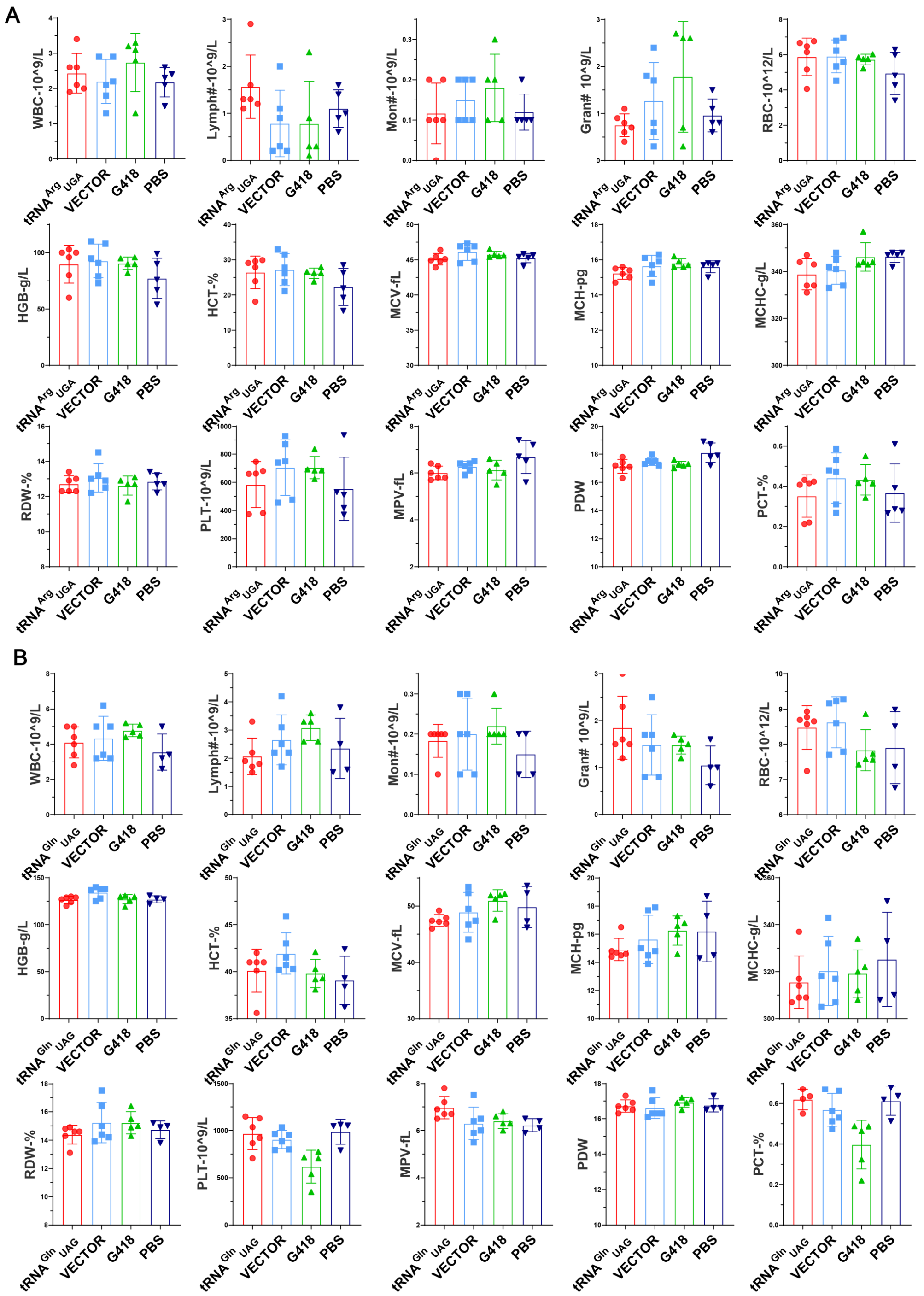


**Figure S20. Blood examination in sup-tRNA in-*vivo* treatment**

(A and B) Safety of the sup-tRNA treatment on Calu-6 (A) and Caco-2 (B) xenograft nude mice assessed by blood routine examination. Data are mean ± s.d. of at least four biological replicates.

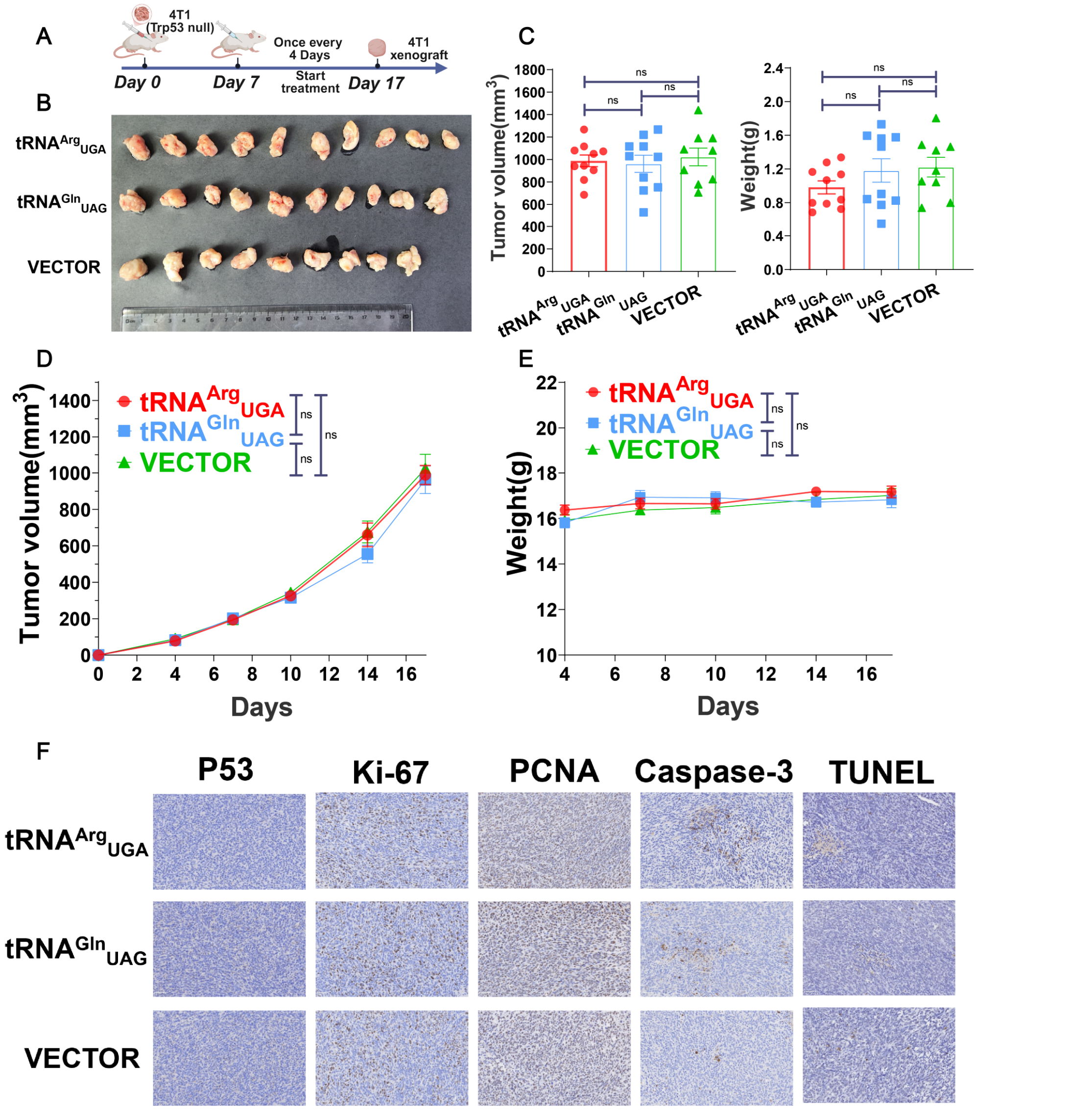


**Figure S21. Therapeutic efficacy of sup-tRNA therapy in *TP53-null* xenograft model**

(A) Schematic of therapeutic evaluation of sup-tRNA therapy in 4T1(*TP53-null*) xenograft model.

(B) Images of 4T1 xenograft excised tumors.

(C) Tumor volumes (left) and weights(right) of tumor collected on in 4T1 xenograft model after sup-tRNA treatment. *P*-values were calculated by one-sided t test.

(D and E) Tumor growth curves(D) and weight curve(E) of 4T1 xenograft model after sup-tRNA treatment. *P*-values were calculated by one-sided t test.

(F) Immunohistochemistry analysis of 4T1 xenograft including p53, proliferation and apoptosis markers after corresponding treatments. Scale bar=50 μm. Data shown here were representative examples from n=3 independent biological replicates.

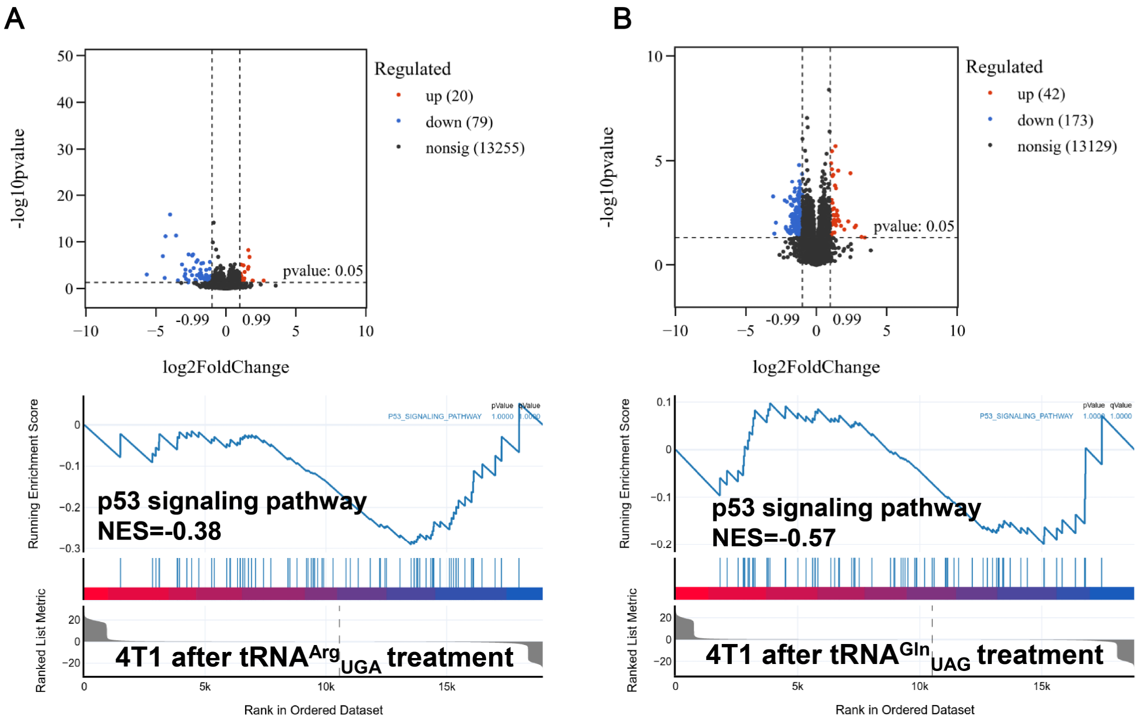


**Figure S22. RNA-seq examination of sup-tRNA in *TP53-null* xenograft model**

(A and B) Gene expression analysis (above) and Gene Set Enrichment Analysis (GSEA) of p53 signaling pathways (below) of 4T1 xenograft treated with tRNA$^{Arg}_{UGA}$(A) and tRNA$^{Gln}_{UAG}$(B).

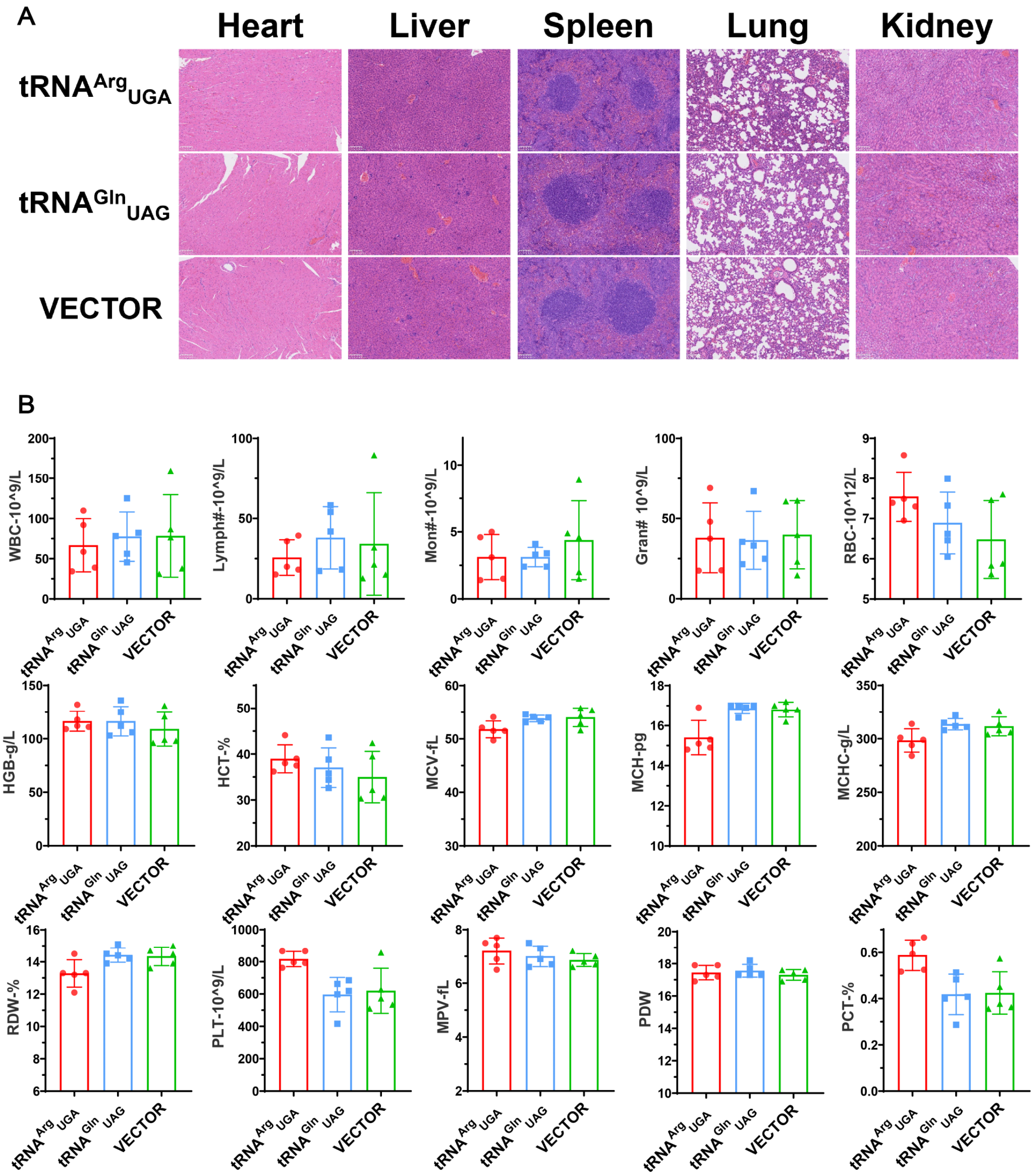


**Figure S23. Safety evaluation of sup-tRNA in *TP53-null* xenograft model**

(A) Safety of the sup-tRNA treatment on heart, liver, spleed, lung, and kidney of 4T1 xenograft nude mice assessed by H&E staining. Scale bars = 100 μm. Data shown here were representative examples from n=3 independent biological replicates.

(B) Safety of the sup-tRNA treatment on 4T1 xenograft nude mice assessed by blood routine examination. Data are mean ± s.d. of at least four biological replicates.
